# CCDC32 collaborates with the membrane to assemble the AP-2 clathrin adaptor complex

**DOI:** 10.1101/2025.08.05.668722

**Authors:** Dillon E. Sloan, Ariel Matthews, Haruaki Yanagisawa, Tanita Tedamrongwanish, Kevin Cannon, Jake Simmons, Garrett Chappell, Nathan I. Nicely, Rebecca Berlow, Masahide Kikkawa, Richard W. Baker

## Abstract

Cells have evolved a variety of assembly chaperones to aid in the difficult process of forming macromolecular complexes in a crowded cytoplasm. Assembly of adaptor protein complex 2 (AP-2), the primary cargo adaptor in clathrin-mediated endocytosis, is regulated by the chaperones AAGAB and CCDC32, whose deletion causes loss of all AP-2 subunits *in vivo*. AAGAB and CCDC32 are thought to act sequentially to assemble the AP-2 tetramer from its constituent heterodimers. However, the molecular requirements and structural consequences of CCDC32 interaction with AP-2 are not yet understood. Here, using *in vitro* reconstitution and integrative structural analysis, we describe the molecular mechanism of CCDC32-mediated AP-2 assembly. First, CCDC32 interacts with the appendage domain of the AP-2 α subunit, using the same binding site as canonical endocytic regulators in addition to a novel, yet highly conserved pocket on α. CCDC32 contains cargo sorting motifs normally found in trans-membrane cargo and binds to AP-2 heterodimers using canonical cargo-binding sites. Additionally, two amphipathic helices in CCDC32 bind to the α/σ2 heterodimer. Surprisingly, in solution, we find that CCDC32 prevents complex assembly and actively disassembles AP-2 tetramers. Inhibition requires the amphipathic helices of CCDC32, which also mediate binding to PIP2-containing membranes. The presence of PIP2-containing membrane stabilizes the final stages of assembly. We propose that the membrane acts as a molecular switch to release inhibitory interactions, allowing for full complex assembly to proceed. Using cryo-EM, we visualize an assembly intermediate that mimics the conformation of AP-2 found in vesicles, with CCDC32 bound at both cargo binding sites and both membrane-binding sites, suggesting that assembly leads to deposition of active complexes on the plasma membrane.

## Introduction

Human cells contain thousands of stable protein complexes^1^, which require the concerted action of molecular chaperones for correct assembly. Molecular chaperones mediate both the folding of individual polypeptides and their subsequent assembly into multi-subunit complexes^2,3^. In general, folding chaperones ensure that the proper *intra-*molecular interactions occur, while assembly chaperones do the same for *inter-*molecular interactions. Assembly chaperones thus assist in appropriate protein complex formation, but they themselves are not a part of the final complex^2^. Many examples of assembly chaperones exist, including those which mediate the formation of the carbon-capture enzyme complex Rubisco, a hexadecamer composed of eight large and small subunits^4^. Other disparate systems have been shown to have dedicated assembly chaperones^5^, like primary cilia assembly^6^ and nucleosome and histone assembly^7^. More recently, a dedicated assembly chaperone system for a subset of the clathrin-adaptor complexes has been described^8,9^.

The adaptor protein complex 2 (AP-2) is the primary cargo adaptor for clathrin-mediated endocytosis^10^. As AP-2 interacts directly with the membrane, cargo, and clathrin, it is often described as the central regulatory hub for endocytosis^11,12^. AP-2 interacts with the plasma-membrane via multiple phosphatidylinositol-(4,5)-bisphosphate (PIP2) binding sites, the strongest of which reside on the α and μ2 subunits. AP-2 also contains two distinct and non-overlapping binding sites for cargo — an “acidic dileucine” or simply “dileucine” cargo binding site on the α/σ2 subunits which recognize trans-membrane proteins with a [D/E]xxxL[L/I] motif, and a “tyrosine” cargo binding site on the μ2 subunit which recognizes cargo bearing a Yxxϕ motif, where ϕ is any bulky hydrophobic residue^13–16^. AP-2 also directly binds clathrin through a clathrin-binding box in the β2 hinge and incorporation of the β2 appendage into the clathrin lattice^17–19^. Notably, all of these motifs are occluded when AP-2 is in the cytosol and only become exposed when AP-2 is on the plasma membrane in an active conformation^12,16,19–21^. Finally, AP-2 contains a number of binding sites for endocytic regulators on the α and β2 appendage domains, which are flexibly tethered to the core of AP-2^22^. The multiplicity of AP-2 binding events and a discrete series of conformational re-arrangements underlie the logic of AP-2 recruitment, activation, and packaging into endocytic structures^23^. AP-2 domain architecture and general interactions are summarized in Fig. 1A.

**Figure 1:**
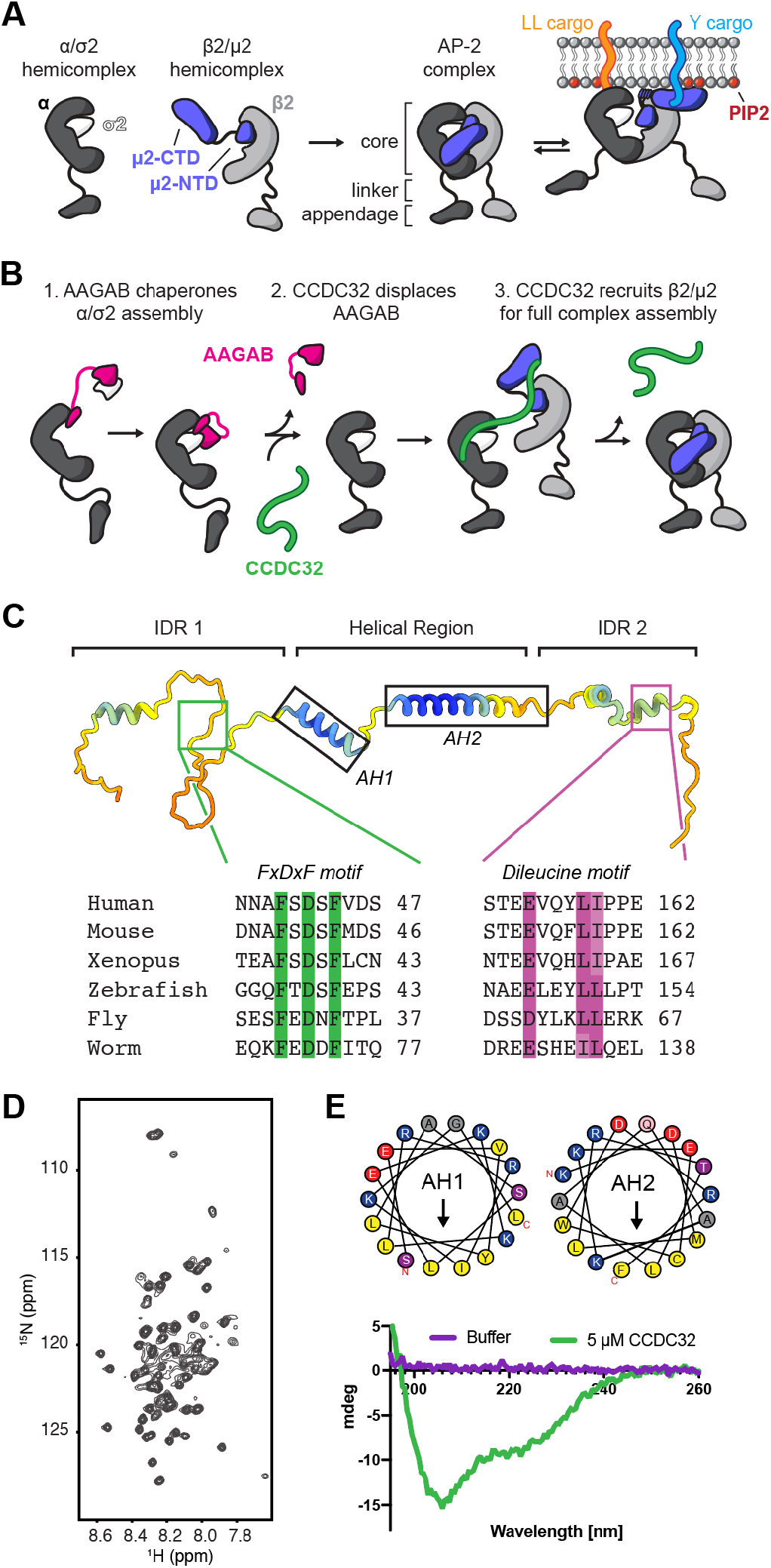
CCDC32 is an intrinsically disordered protein with canonical AP-2 binding motifs. **(A)** Schematic showing the domain organization of AP-2 core and the two hemicomplexes. AP-2 undergoes a conformational change at the membrane, allowing for cargo binding. LL = dileucine cargo motif, Y = tyrosine cargo motif. **(B)** Schematic showing the AP-2 assembly pathway, as mediated by AAGAB and CCDC32. **(C)** AlphaFold2 structure of CCDC32 (Uniprot ID Q8BS39) colored by pLDDT score. Domains are labeled. Canonical AP-2 binding motifs are highlighted and a sequence alignment for standard model organisms is shown. **(D)** ^1^H-^15^N HSQC spectrum of ^15^N-labeled CCDC32. **(E)** A circular dichroism (CD) spectra shows that CCDC32 is partially helical, with a predicted helicity of 44% (bottom). Helical wheel diagram of the two predicted amphipathic helices in CCDC32 (top).

Whereas the function and regulation of AP-2 during endocytosis is well established^23^, assembly of multi-subunit adaptor complexes is comparatively less studied. AP-2 is composed of two large subunits (α and β2), a medium subunit (μ2), and a small subunit (σ2) that assemble hierarchically from hemi-complexes (α/σ2 and β2/μ2) before forming the final hetero-tetramer (Fig. 1A,B). The assembly of AP-2 is regulated by the chaperones AAGAB (<u>a</u>lpha <u>a</u>nd <u>g</u>amma <u>a</u>daptin <u>b</u>inding protein) and CCDC32 (<u>c</u>oiled-<u>c</u>oil <u>d</u>omain-<u>c</u>ontaining protein <u>32</u>, also known as c15orf57)^8,9^ (Fig. 1B). Mutations in AAGAB cause Palmoplantar keratoderma type IA^24,25^, an autosomal dominant hereditary skin disease. AAGAB deletion in cultured cells significantly decreases the amount of all four AP-2 subunits, although it only binds the α/σ2 hemi-complex^8^. Given AAGAB only directly interacts with half of the AP-2 complex and yet is necessary for maintaining levels of all four subunits, it is thought to function as an assembly chaperone for AP-2, as well as for the related complexes AP-1 and AP-4 ^8,26,27^ (Fig. 1B).

Mutations in the gene encoding CCDC32 have been linked to Cardiofacioneurodevelopmental syndrome, an autosomal recessive craniofacial syndrome marked by several developmental abnormalities^28–30^. Animal studies and cellular data suggest a role for CCDC32 in ciliogenesis^28^ and endocytosis^31^, likely dependent on an ability to interact physically with AP-2^32^. Like AAGAB, loss of CCDC32 significantly decreases the steady state level of all four AP-2 subunits in HeLa cells, and CCDC32 can interact with both AP-2 hemicomplexes^9^. Unlike AAGAB, CCDC32 does not appear to interact with other AP complexes. *In vitro*, CCDC32 is able to displace AAGAB from the α/σ2 hemicomplex^9^. After AAGAB displacement, CCDC32 is thought to recruit β2/μ2, which ultimately leads to CCDC32 release and the completion of AP-2 assembly^9^ (Fig. 1B). The molecular requirements and structural determinants of this process have not yet been determined.

In this study, we have used structural, biochemical, and bioinformatic methods to determine the mechanism of CCDC32-mediated AP-2 assembly. CCDC32 first encounters AP-2 via the α appendage domain using a novel binding motif not present in other endocytic proteins. CCDC32 contains two other binding sites for the α/σ2 core, a canonical dileucine cargo motif and an α helical domain that mimics an intramolecular interaction in the α subunit. We show that AAGAB displacement is mediated by direct competition for the dileucine binding pocket on AP-2 σ2. Subsequent recruitment of the β2/μ2 hemicomplex is mediated by a direct interaction with the μ2 C-terminal domain (CTD), in a multivalent fashion that requires interaction with both the PIP2 and cargo-binding sites on μ2. In this manner, CCDC32 utilizes two cargo binding sites and membrane binding sites to mediate complex assembly. Using cryo-EM, we visualize two assembly intermediates, showing that recruitment of β2/μ2 displaces CCDC32 motifs required for α binding, thereby describing the stepwise interaction network that underlies the hemicomplex to assembled complex transition. Surprisingly, CCDC32 acts as an assembly inhibitor in a soluble environment. We find that CCDC32 is a *bona fide* membrane-binding protein that uses the same alpha helical domains that mediate binding to the α/σ2 assembly intermediate as it does to bind to the membrane. The membrane therefore acts as a switch to release inhibitory binding interactions, allowing full assembly to occur. We show that addition of PIP2 containing membranes in the presence of CCDC32 stabilizes the assembled AP-2 tetramer, suggesting a potential role for the plasma membrane in ultimate AP-2 assembly.

## Results

### CCDC32 contains canonical AP-2 endocytic motifs

While CCDC32 has been implicated as an assembly factor for AP-2, its mechanism of action has not been reported. To understand this mechanism, we set out to find the molecular determinants and structural consequences of the CCDC32/AP-2 interaction. CCDC32 is a small ∼20 kDa protein that is predicted by AlphaFold^33^ and Metapredict^34^ to be an intrinsically disordered protein, with N- and C-terminal intrinsically disordered regions that flank a central partially helical domain (Figure 1C, Supp. Fig. 1A,B). The ^1^H-^15^N HSQC NMR spectrum of ^15^N-labeled CCDC32 confirms that CCDC32 is largely disordered, with all of the observable crosspeaks appearing in a narrow ^1^H chemical shift range that is characteristic of an intrinsically disordered protein (Fig 1D, Supp. Fig. 1C). The central region of the protein is predicted to contain two conserved alpha helices, which appear to be amphipathic in nature based on their high hydrophobic moments and helical wheel plots (Figure 1C,E; Supp. Fig. 1D). This was confirmed with circular dichroism (CD) spectroscopy, which showed ∼44% helical content using fulllength mouse CCDC32, in agreement with the amount of secondary structure predicted by AlphaFold (Figure 1E). We refer to these amphipathic alpha helices as AH1 and AH2.

We next used bioinformatic characterization to assess conservation and look for known AP-2 binding motifs in CCDC32. AP-2 σ2 is conserved in most eukaryotes with sequenced and annotated genomes, showing AP-2 is broadly conserved (Supp. Fig. 1E). AAGAB is largely conserved in metazoa, plants, and fungi, while CCDC32 is conserved exclusively in metazoa (Supp. Fig. 1E). Analysis of CCDC32 sequence conservation using Consurf^35^ (Supp. Fig. 2A) revealed several predicted functional regions within the protein, including three potential canonical AP-2 binding motifs: a DLW motif at aa17-19, an FxDxF motif at aa39-43 and a dileucine cargo motif at aa154-159 (Figure 1C; Supp. Fig 2B-H; amino acid position reflects the *Mus musculus* ortholog sequence unless otherwise noted). These putative AP-2 binding motifs are conserved in most CCDC32 orthologs in both sequence and general position in the protein (Supp. Fig. 2I,J). Importantly, none of the CCDC32 orthologs identified were predicted to be integral membrane proteins, unlike traditional AP-2 cargo proteins that bear a dileucine motif. Vertebrate orthologs of CCDC32 contain a Wxxϕ motif at aa68-71 (Supp. Fig. 2D), recently identified in FCHo2 as mediating binding to the μ2 CTD by mimicking the tyrosine-based cargo motif Yxxϕ^36^. AH1 and AH2 are both highly conserved (Supp. Fig. 2F,G). The intrinsically disordered regions are also enriched in acidic residues, glycines, and prolines, while the central AH1 and AH2 domains are enriched in basic residues (Supp. Fig. 2I). Together, this data suggests that the mode of interaction of CCDC32 and AP2 is largely conserved.

**Figure 2:**
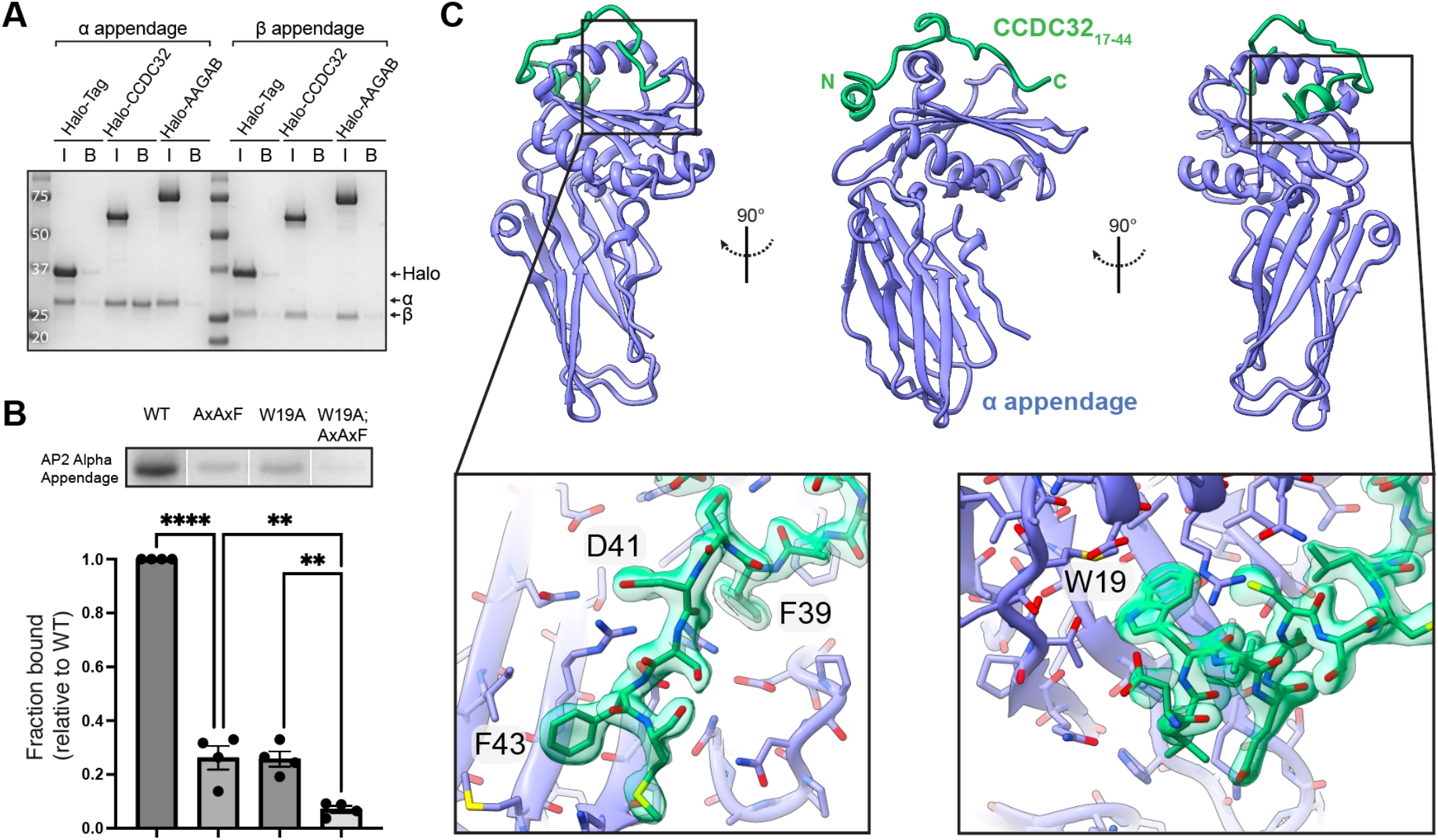
CCDC32 binds to the α appendage via a non-canonical extended FxDxF motif. **(A)** Pull-down binding assay with Halo-tagged CCDC32 and AAGAB with the α and β2 appendages. Bound proteins were eluted by boiling the beads, as cleaved CCDC32 and the α appendage otherwise co-migrate as a single band on an SDS-PAGE gel. HaloTag-CCDC32 and AAGAB are not in the elution lanes because they are covalently linked to the resin via the Halo tag. I = Input; B = Bound. **(B)** Pull-down binding assay with Halo-tagged CCDC32 and various mutants with the α appendage. The full, uncropped gel is labeled and shown in Supp. Fig. 3A. Error bars represent the standard error of the mean of four replicates. ** = p<0.01; **** = p<0.0001. **(C)** X-ray crystal structure of the α appendage in complex with aa17-44 of CCDC32, shown in three views. Zoom-in panels show the electron density for the FxDxF motif (left) and W19 (right).

### CCDC32 interacts with AP-2 α appendage via a novel, extended FxDxF motif

The α appendage is a hotspot for AP-2 recruitment, interacting with a number of small peptide motifs (FxDxF, D[PL][FW], Wxx[FW]x[DE]) that are contained within many endocytic regulators, like EPS15, Epsin, and Amphiphysin^37^. CCDC32 is predicted to contain at least two appendage binding motif sequences, a DLW motif and an FxDxF motif, the latter of which (FSDSF) matches the optimal binding sequence^22^. Using Halo-tagged CCDC32 in a pull-down assay, we show that CCDC32 indeed binds to the α appendage, but not the β2 appendage (Figure 2A). AAGAB does not bind to either appendage (Figure 2A). Mutation of FxDxF to AxAxF decreased, but did not eliminate binding, as did mutation of DLW to DLA (Figure 2B; Supp. Fig. 3A). Mutating both sites effectively eliminates α appendage binding to CCDC32 (Figure 2B; Supp. Fig. 3A). The relative position of the DLW and FxDxF motifs in CCDC32 with respect to each other appears to be tightly controlled across evolution (Supp. Fig. 2J). However, the spacing of the known binding sites for DLW and FxDxF motifs on the α appendage would preclude simultaneous binding of both motifs. We used AlphaFold3^38^ to model the configuration of these motifs in binding the α appendage, which showed with high confidence that DLW binds to a distinct, highly conserved part of the α appendage not identified in previous structures (Supp. Fig. 3B-D). To determine the molecular details of this interaction, we used a 28 amino acid peptide spanning both sequence motifs and determined an X-ray crystal structure of CCDC32 bound to the α appendage (Figure 2C, Table 1). This shows that CCDC32 binds to the FxDxF binding site in a nearly identical fashion to other FxDxF motif-containing proteins, but the N-terminal residues make extensive contact with the top of the appendage and bury the highly conserved W19 in a hydrophobic pocket on the backside of the appendage (Figure 2C; Supp. Fig. 3C,D; Supp. Fig. 4A,B). We also determined a crystal structure of only the CCDC32 FxDxF motif bound to the α appendage (Table 1). While this shows a nearly identical structure at the FxDxF motif binding pocket, it clearly shows that W19 binding results in a hydrophobic pocket closing around CCDC32 (Supp. Fig. 4B). This displaces 6 water molecules and causes a 6 Å displacement of a loop in the appendage (Supp. Fig. 4B). Fluorescence polarization estimated that the extended motif binds with 287 nM affinity (Supp. Fig 4C), an order of magnitude tighter binding compared to any other FxDxF or D[PL][FW] motif in isolation^22^. As expected, the FxDxF of CCDC32 alone has a Kd of 4.15 μM, in line with other FxDxF motifs (Supp. Fig 4C). How such a high-affinity interaction is displaced after AP-2 assembly concludes is an open question.

**Table 1.** X-ray collection and refinement statistics.

|  | AP2 alpha appendage + CCDC32 FxDxF motif | AP2 alpha appendage + CCDC32 extended FxDxF motif |
| --- | --- | --- |
| **PDB ID** | 9PUE | 9PPP |
| **Data reduction** |  |  |
| Spacegroup | P<sub>21</sub> | R<sub>3</sub> |
| Cell |  |  |
| a, b, c (Å) | 40.807, 72.465, 42.053 | 96.664, 96.664, 195.231 |
| α, β, γ (°) | 90 99.092 90 | 90 90 120 |
| Resolution range (Å)* | 21.59-4.10 (1.97-1.91) | 24.17-2.10 (2.18-2.10) |
| Number total/unique reflections | 179063/18829 (15831/1771) | 541715/39669 (34758/3930) |
| Multiplicity | 9.5 (8.9) | 13.7 (8.8) |
| Completeness (%) | 99.6 (95.9) | 100 (100) |
| I/sig(I) | 10.1 (2.4) | 16.1 (2.8) |
| R<sub>merge</sub> (%) | 18.6 (86.2) | 15.0 (80.1) |
| R<sub>pim</sub> (%) | 6.3 (30.1) | 4.2 (28.3) |
| CC<sub>1/2</sub> | 99.5 (85.5) | 99.8 (86.3) |
| **Structure refinement** |  |  |
| Resolution (Å) | 20.88-1.91 (1.93-1.91) | 23.43-2.10 (2.13-2.10) |
| R<sub>work</sub>/R<sub>free</sub> | 18.23/23.02 (23.49/24.87) | 16.72/20.41 (26.27/27.61) |
| Number atoms | 2212 | 4877 |
| RMSD* |  |  |
| Lengths (Å) | 0.007 | 0.008 |
| Angles (°) | 0.805 | 0.893 |
| B factors |  |  |
| Avg. B all atoms (Å<sup>2</sup>) | 19.01 | 25.75 |
| Avg. B protein atoms (Å<sup>2</sup>) | 18.54 | 24.77 |
| Avg. B ligand atoms (Å<sup>2</sup>) | 43.66 | 34.66 |
| Avg. B water molecules (Å<sup>2</sup>) | 24.23 | 32.50 |
| Ramachandran plot |  |  |
| Favored (%) | 97.99 | 97.34 |
| Allowed (%) | 2.01 | 2.66 |
| Outlying (%) | 0 | 0 |
| Clashscore | 2.48 | 3.51 |

**Figure 3:**
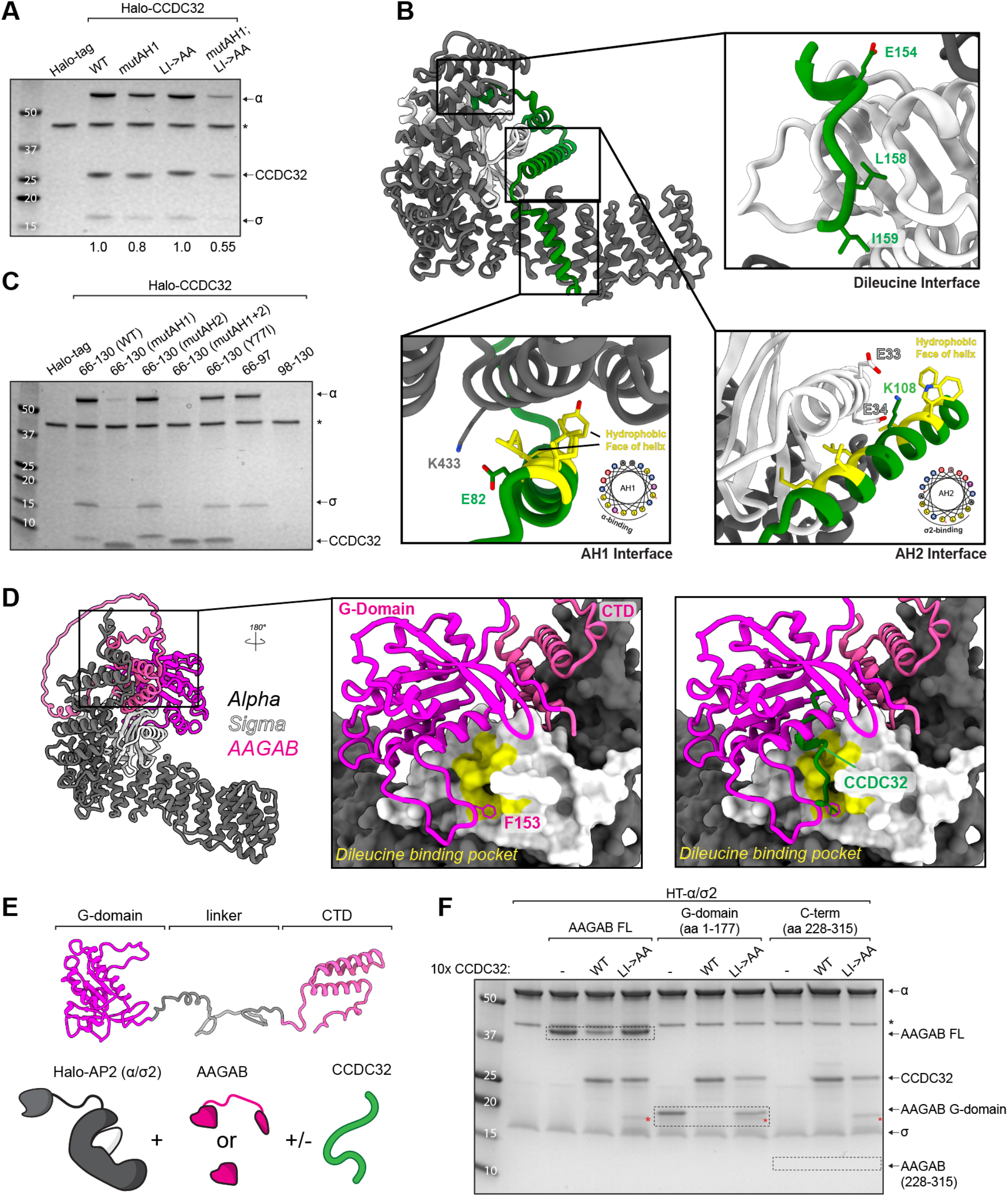
CCDC32 binds to the α/σ2 hemicomplex in a multivalent fashion and displaces AAGAB via its dileucine motif. **(A)** Halo-tag pulldown binding assay between HT-CCDC32 and the α/σ2 hemicomplex. Mutation of AH1 and the dileucine motif reduce binding and nearly abolish binding in combination. mutAH1 = L78A/L81A/K83A/K84. * denotes HRV, used to elute complexes off the Halo resin. Numbers below gel lanes represent α/CCDC32 band intensity ratio, normalized to WT. **(B)** AlphaFold3 prediction of α/σ2 (grey/white) with CCDC32 (green). Binding of AH1 on α is circled in red and binding to the dileucine pocket on σ2 is shown in inset. **(C)** Halo-tag pulldown binding assay between HT-CCDC32 AH1+AH2 (aa66-130) and the α/σ2 hemicomplex. AH1 is necessary and sufficient for binding. A CCDC32 Y77I mutant does not break binding. mutAH1 = L78A/L81A/K83A/K84A. mutAH2 = V95S/M100S/L101S/L104S. mutAH1 runs at a lower MW because multiple charged residues are mutated to alanine. * as in (**A**). **(D)** AlphaFold3 prediction of α/σ2 (grey/white) with FL AAGAB (G-domain: magenta; CTD:pink). Binding to the dileucine pocket on σ2 (yellow) is shown in inset, with surface representation for AP-2 subunits. CCDC32 from another Alphafold model is shown in green, showing an overlap at the dileucine binding pocket. **(E)** Schematic for a competition binding assay between HT-α/σ2, CCDC32, and AAGAB. **(F)** Competition binding assay showing that CCDC32 outcompetes AAGAB for α/σ2 binding at the dileucine cargo binding pocket. Red * denotes a contaminant at a similar size as AAGAB 1-177. Dashed boxes highlight the various AAGAB constructs used. An input gel is shown in Supp. Fig. 6C.

**Figure 4:**
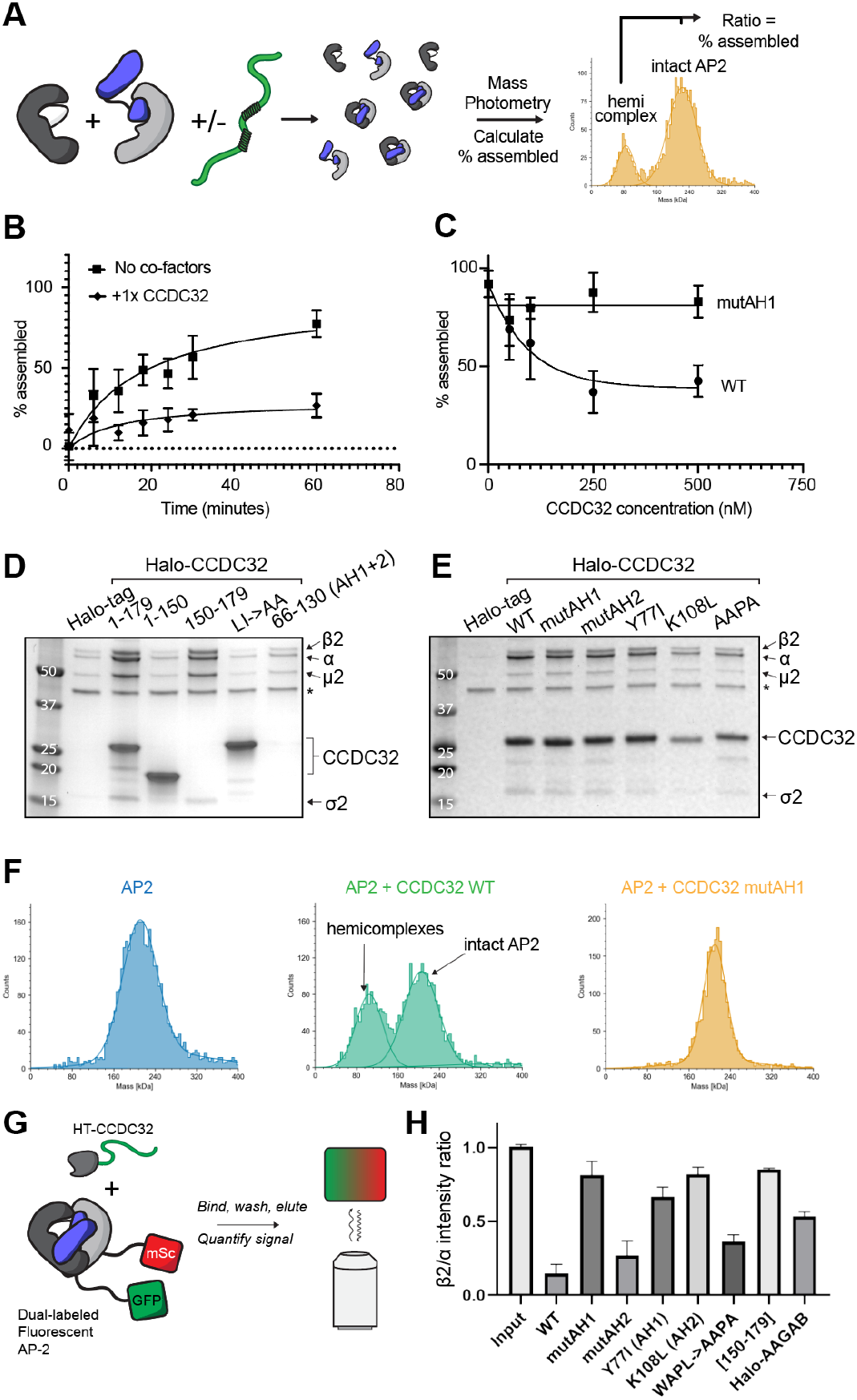
CCDC32 regulates AP-2 (dis)assembly via AH1 and AH2. **(A)** AP-2 assembly assay via mass photometry. Hemicomplexes are incubated for various times +/-CCDC32, and assembly is measured as the ratio of peaks corresponding to hemicomplex or full complex. **(B)** Concentration dependence of AP-2 assembly. **(C)** Time-dependent assembly assay shows that 1x CCDC32 acts as an assembly inhibitor, where 1/10x CCDC32 has no effect. **(D)** Halo-tag pulldown binding assay between HT-CCDC32 constructs and AP-2 core. Note the ratio of the α and β2 subunit between conditions. * denotes HRV, used to elute complexes off the Halo resin. **(E)** Halo-tag pulldown binding assay between HT-CCDC32 constructs and AP-2 core. Note the ratio of the α and β2 subunit between conditions. mutAH1 = L78A/L81A/K83A/K84. mutAH2 = V95S/M100S/L101S/L104S. * as in (**A**). **(F)** Mass photometry size distribution profiles of AP-2, AP-2 co-incubated with CCDC32, and AP-2 co-incubated with mutAH1 CCDC32. **(G)** Schematic of dual-labeled fluorescent AP-2 pull-down assay. α is tagged with GFP and β2 is tagged with mScarlet. The ratio of the fluorescence channels is measured in a plate reader after a Halo-tag pull-down assay with HT-CCDC32. **(H)** Bar graph of β2/α fluorescence intensity ratios after a Halo-tag pulldown, showing the mean and standard deviation of three replicates. All readings are normalized to an input of β2/α. Note: this graph does not show differences in affinity between the CCDC32 constructs, only the ratio of α/β2. Relative affinity differences can be seen in panel **D** and **E**.

To see if other proteins contain this extended FxDxF motif, we first compared all reported trafficking proteins bearing an FxDxF motif (Supp. Fig. 4D). Only CD2AP and PLEKHM2 have a tryptophan in the first 20 amino acids preceding the FxDxF motif, but neither show an interaction with the W-binding pocket when modeled using AlphaFold3. To take a more systematic approach, we looked for FxDxF sequences in all annotated ORFs in Uniprot from the mouse proteome, yielding 922 unique sequences. Of these, only 62 had a tryptophan in the 30 amino acids preceding the FxDxF motif. None had a tryptophan in the optimal distance of 17-21 amino acids. After removing sequences in folded regions, AlphaFold3 modeling did not show interaction for any candidates in the same manner as CCDC32. While other proteins may bind to this newly identified conserved binding pocket on the α appendage, we were unable to find additional proteins with the same extended FxDxF motif as CCDC32.

### CCDC32 interacts with AP-2 α/σ2 core via its dileucine motif and amphipathic helix 1 (AH1)

CCDC32 has previously been shown to bind the α/σ2/AAGAB ternary complex, although the binding motifs on CCDC32 mediating this interaction are unknown^9,32^. Our bioinformatic approach suggests that the α/σ2 interaction is likely mediated by the dileucine motif in CCDC32. Using Halo-tagged CCDC32 in a pull-down assay, we were able to show that Halo-CCDC32 robustly binds recombinant α/σ2 (Fig. 3A). Reciprocally, Halo-α/σ2 is able to pulldown CCDC32 (Supp. Fig. 4E). Halo-α/σ2 is also able to pull down the α appendage, but only in the presence of CCDC32, showing that the same molecule of CCDC32 is able to simultaneously engage the α/σ2 core and the α appendage (Supp. Fig. 4E,F).

Unexpectedly, mutation of the dileucine motif (LI->AA) did not seem to affect binding of CCDC32 to α/σ2 (Fig. 3A). Using multiple structure prediction algorithms to model the interaction of CCDC32 with the α/σ2 core, we find three potential interaction sites for CCDC32 (Figure 3B; Supp. Fig. 5A-C). These same interfaces are generally predicted for other CCDC32 orthologs, as well (Supp. Fig. 5D). First, the dileucine motif is predicted to bind to the dileucine binding site on σ2. Next, both AH1 and AH2 are predicted to make direct interaction, with AH1 packing against α and AH2 packing against a side of σ2 distal from the dileucine binding site (Fig. 3B; Supp. Fig. 5A-B). While both AH1 and AH2 are highly conserved at the amino acid level (Supp. Fig. 2A,F,G), AH1 was modeled with high confidence in all methods tested, whereas AH2 was only modeled by AlphaFold3 as packing against σ2. While both helices are predicted to bind using their hydrophobic face, each contains a charged residue that is in the vicinity of complementary residues on α/σ2, potentially lending specificity to the interaction (Fig. 3B, inset; Supp. Fig. 5B, inset). While the local pLDDT scores are high (generally >80), and the PAE matrix plot shows confidence in all predicted interfaces (Supp. Fig. 5B,C), it is important to emphasize that the exact molecular nature of these interfaces is still uncertain.

**Figure 5:**
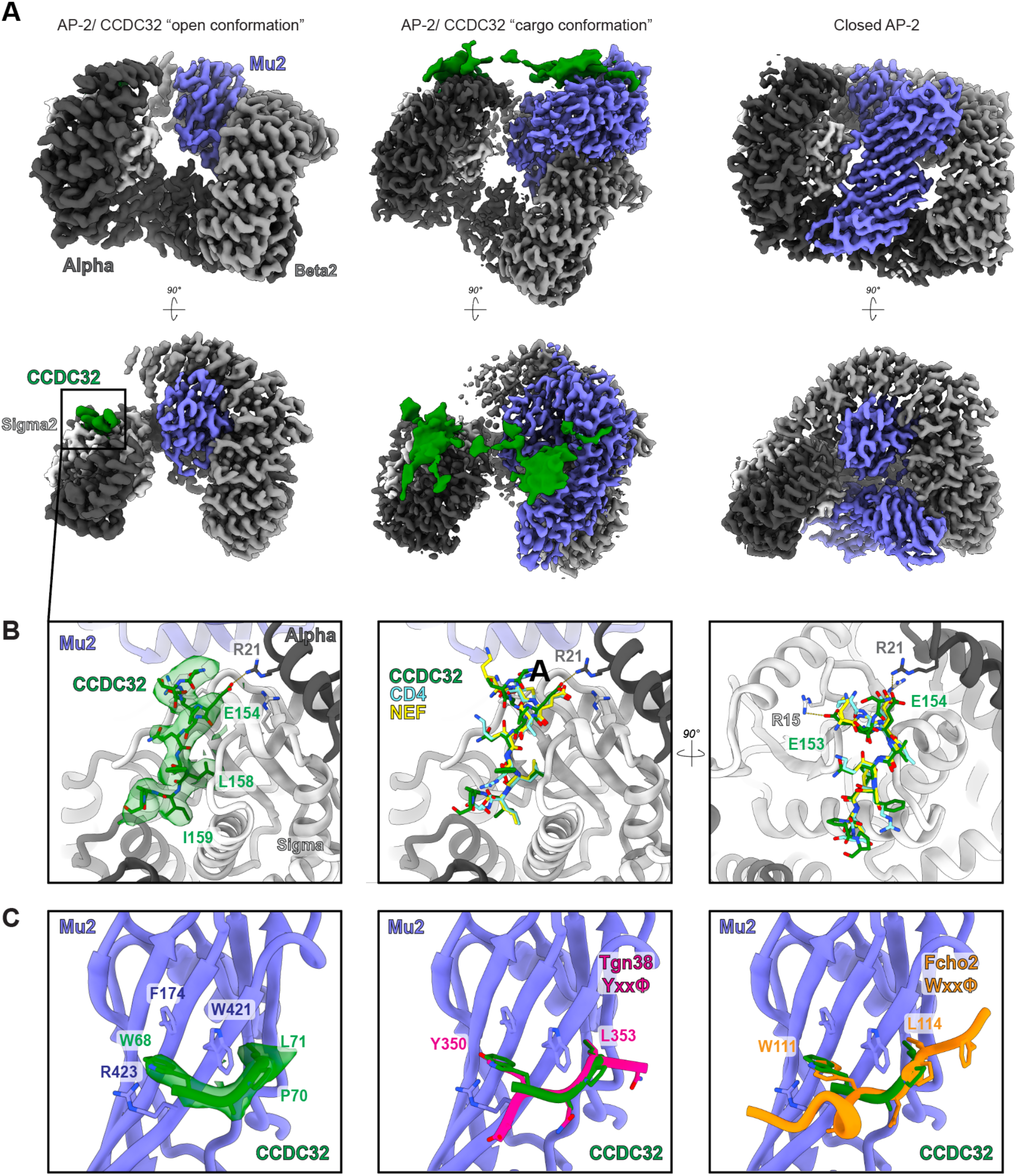
Cryo-EM reveals intermediates of CCDC32-mediated AP-2 assembly. **(A)** Three cryo-EM structures of AP-2 determined after co-incubation of AP-2 with CCDC32. Left: “Bowl” AP-2/CCDC32 where CCDC32 is only observed at the dileucine binding pocket and the μ2-CTD is not in a fixed location. Middle: “cargo-bound” AP-2/CCDC32 conformation, where AP-2 adopts the conformation seen on membranes. The μ2-CTD is in a fixed location and density for CCDC32 is observed on the dileucine cargo binding pocket, the tyrosine cargo binding pocket, and both PIP2 binding sites. Right: closed AP-2, i.e. the assembly product, is observed with no density for CCDC32. **(B)** Zoom-in view of the dileucine cargo binding site in the “bowl” structure. CCDC32 adopts a canonical dileucine binding mode, when compared to CD4 and NEF. Right: CCDC32 has an additional salt bridge mediated by a glutamate outside of the ExxxLI motif. **(C)** Zoom-in view of the tyrosine cargo binding site in the “cargo” structure. CCDC32 uses a non-canonical WAPL (Wxxϕ) motif to bind in the same location as other tyrosine-containing cargo, like Tgn38 (magenta). CCDC32 WAPL binding is nearly identical to Fcho2 Wxxϕ binding (orange).

Interestingly, AH1 is predicted to bind in the same binding site as a C-terminal helix of α, which packs against itself when AP-2 is in the open conformation (we refer to this as the “alpha tail”) (Supp. Fig. 6A,B). Pull-down assays using AH1+AH2 (CCDC32^66-130^), AH1 alone (CCDC32^66-97^), or AH2 alone (CCDC32^98-130^) show that AH1, but not AH2, is sufficient for binding α/σ2 core (Fig. 3C). Mutating AH1 to L78A/L81A/K83A/K84A (CCDC32^66-130;mutAH1^) prevents binding of the truncation (Fig. 3C). This suggests that the interface predicted from the Alphafold modeling is indeed directly contacting α/σ2. Mutating both AH1 and the dileucine motif in full-length CCDC32 significantly reduces binding to AP-2 α/σ2 core (Figure 3A). Overall, this shows that CCDC32 binds to α/σ2 in a multivalent manner, using at least three (extended FxDxF, dileucine, AH1) and possibly four (AH2) binding sites. To our knowledge, this is the first endogenous regulator described as binding to the dileucine cargo binding site on AP-2, excluding the HIV protein Nef^39^. CCDC32 is believed to displace AAGAB from the α/σ2 hemicomplex during AP-2 assembly^9^. Given CCDC32 contains a dileucine cargo motif, and mutating the dileucine binding site in AP-2 α/σ2 reduces binding of AAGAB^8^, we hypothesized interaction of CCDC32’s dileucine motif with the AP-2 α/σ2 core to be important for AP-2 assembly by displacing AAGAB. AlphaFold modeling of the interaction of AAGAB with α/σ2 suggests that the N-terminal domain of AAGAB (G protein-like, or G-domain) binds via the dileucine binding pocket (Figure 3D), corroborating earlier experiments where mutating the dileucine binding pocket reduces binding of AAGAB^8^. AAGAB F153 is predicted to occupy the dileucine binding pocket, reminiscent of how F7 from β2 packs into this pocket in the closed AP-2 conformation^12^. To test whether CCDC32 competes for binding with AAGAB via the dileucine motif, we used HaloTag pull-down assays with Halo-α/σ2 as bait, AAGAB as prey, and WT and mutant CCDC32 added *in trans* at a 10x molar excess (Fig. 3E). We found that while CCDC32 only moderately outcompeted full-length AAGAB for binding α/σ2, it fully outcompeted binding of the AAGAB G-domain (Figure 3F, Supp. Fig. 6C). An LI->AA mutant of CCDC32 was unable to outcompete AAGAB. The C-terminal domain of AAGAB did not show appreciable binding (Figure 3F). Overall, our data show the multivalent interaction of CCDC32 with the α/σ2 core and provide a mechanistic basic for a core function of CCDC32, which is displacement of the assembly chaperone AAGAB.

**Figure 6:**
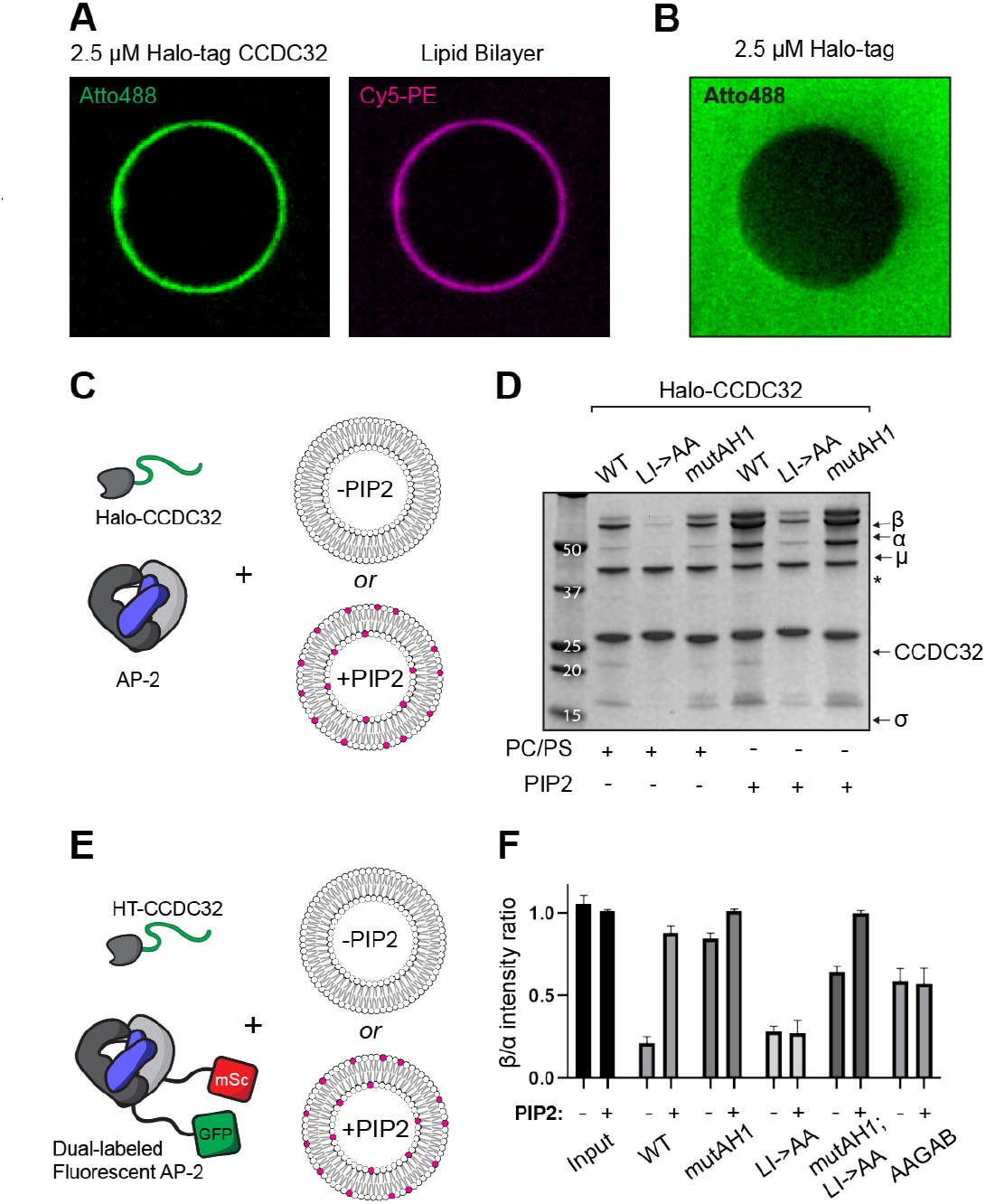
PIP2-containing membranes stabilize AP-2 assembly. **(A)** Membrane-binding assay showing Atto488-labeled Halo-CCDC32 binding to supported lipid bilayers (SLBs). Membrane composition is 74% PC/ 20% PS/ 5% PIP2/ 1% Cy5-PE. Lipid bilayer is imaged using fluorescent lipid (Cy5-PE). **(B)** Membrane-binding assay showing that Atto488-labeled Halo-tag does not bind to the same membrane composition used in (**A**). **(C)** Schematic for Halo-tag pulldown binding assay in the presence of liposomes containing either PC/PS or PC/PS + 5% PIP2. **(D)** Halo-tag pulldown binding assay of various CCDC32 constructs and AP-2 core in the presence of membranes. mutAH1 = L78A/L81A/K83A/K84. Overall, the presence of membranes stimulates CCDC32/AP-2 binding and stabilizes equimolar α/β2 ratios. **(E)** Schematic for Halo-tag pulldown binding assay in the presence of liposomes containing either PC/PS or PC/PS + 5% PIP2 using dual-labeled fluorescent AP-2. **(F)** Halo-tag pulldown binding assay of various CCDC32 constructs and dual-labeled fluorescent AP-2 core in the presence of membranes. Overall, the presence of membranes stabilizes equimolar α/β2 ratios. AAGAB is included, showing this phenomenon is unique to CCDC32.

### CCDC32 binds the μ2 CTD in a multivalent manner

CCDC32 is thought to complete the assembly of AP-2 in the cytosol by recruiting the AP-2 β2/μ2 hemi-complex and subsequently dissociating from the assembled complex^9^ (Fig. 1B). Co-IP experiments showed that CCDC32 directly binds μ2, but not β2^9^. We reasoned that CCDC32 would likely interact with the μ2-CTD, as the N-terminal longin domain is largely occluded in the hemicomplex and assembled AP-2. Using a Halo-tag pulldown assay, we found that CCDC32 directly binds the μ2-CTD (Supp. Fig. 7A). Using truncations of CCDC32 (aa 1-100 and aa 101-179), we identified binding sites for the μ2-CTD at both the N- and C-terminus of the protein (Supp. Fig. 7B). As a construct containing only the AH1 and AH2 domains (a66-130) does not bind the μ2-CTD (Supp. Fig. 7A), the interaction is likely mediated by the intrinsically disordered regions of CCDC32. Further truncations using 30 aa windows did not show binding above background, outside of aa 121-150, indicating that the overall interaction is likely driven through multivalent binding (Supp. Fig. 7B). Interestingly, addition of inositol hexaphosphate (IP6), a soluble analog of the PIP2 lipid headgroup, eliminated binding of CCDC32 (Supp. Fig. 7A). Conversely, a mutant μ2-CTD that prevents binding to PIP2 was unable to bind CCDC32 (Supp. Fig. 7C). Addition of IP6 also prevented binding of the recombinant β2/μ2 core hemicomplex, indicating the primary binding site of CCDC32 to the β2/μ2 hemicomplex is the CTD of μ2 (Supp. Fig. 7D). This IP6-dependent binding of CCDC32 to the β2/μ2 hemicomplex was conserved in 4 out of 5 species of CCDC32 tested in pull-down assays (Supp. Fig. 7D). Therefore, recruitment of the β2/μ2 hemicomplex depends on multivalent interactions with important regulatory sites on the μ2-CTD.

**Figure 7:**
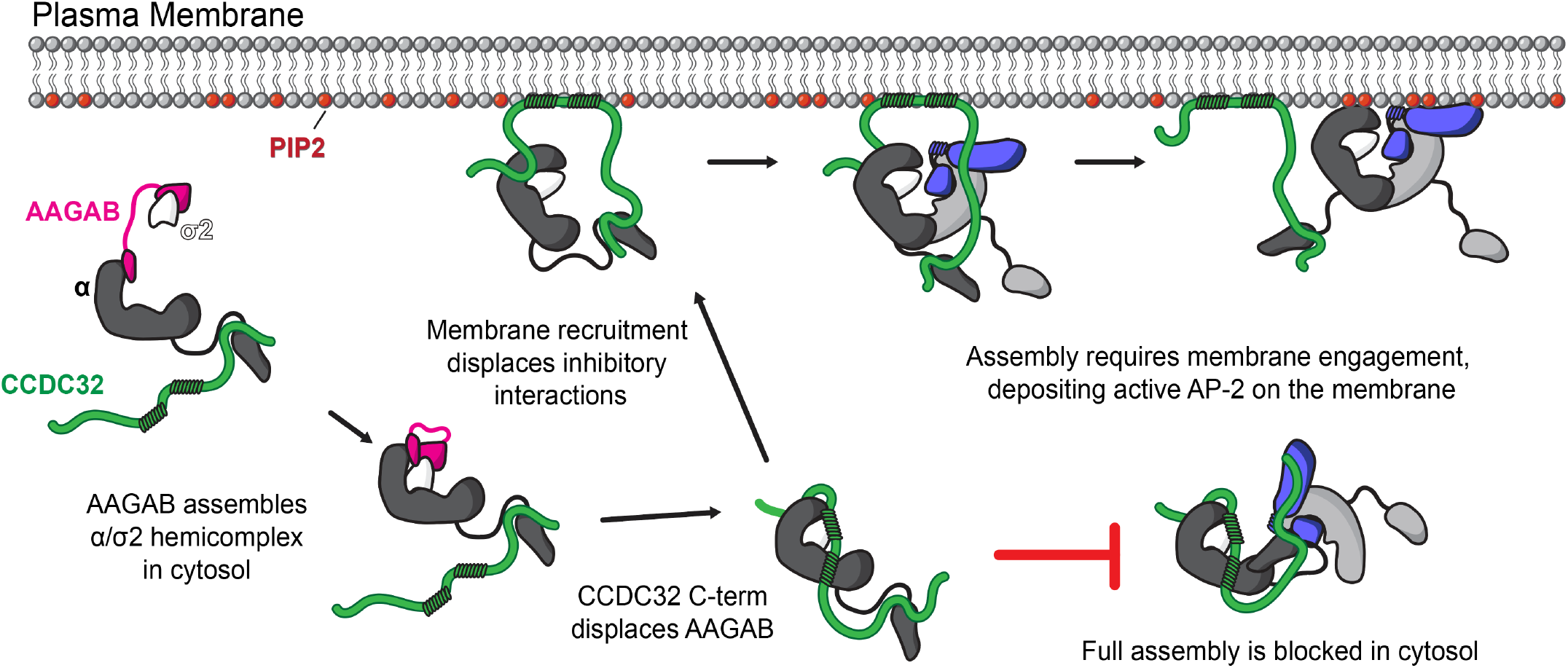
Structure-informed model for CCDC32-mediated AP-2 assembly. Initial α/σ2 complex formation is templated using the two domains of AAGAB, which bind to α and σ2 independently. CCDC32 is able to bind regardless of AAGAB presence via its extended FxDxF motif. After α/σ2 assembly, local concentration effects of CCDC32-appendage tethering drive displacement of AAGAB, which requires the dileucine motif of CCDC32. CCDC32 then binds with its AH1 and AH2 domains, stabilizing the ternary complex. Full assembly of AP-2 is inhibited in the cytosol, where interactions of α/σ2 with AH1/AH2 inhibit β2/μ2 binding. These inhibitory interactions are released at the membrane, where AH domains contact the membrane. Functional AP-2 is not formed until recruitment of β2/μ2 which contains the clathrin binding box. CCDC32 may be displaced via interaction with cargo and other AP-2 regulators, or else may persist and regulate AP-2 via a separate endocytic function.

### CCDC32 inhibits AP-2 assembly in the soluble state

Our analysis to this point yielded a testable mechanism — CCDC32 interacts with both hemicomplexes in a multivalent manner to assemble the full heterotetrameric complex. As CCDC32 is predicted to be an assembly factor for AP-2, the presence of CCDC32 should meaningfully alter the assembly of AP-2 from the component α/σ2 and β2/μ2 hemicomplexes, dependent on its ability to interact using the molecular interfaces we have identified. To test this, we purified hemicomplexes and tested their assembly into AP-2 heterotetramers using mass photometry (Fig. 4A; Supp. Fig. 8A). To test the concentration dependence of AP-2 assembly, we incubated hemicomplexes at various concentrations, then measured the mass distribution profile of the sample using mass photometry. As shown in Supp. Fig. 8B, AP-2 assembly shows a concentration-dependent assembly profile, with AP-2 assembling at both a faster rate and to a higher overall level at higher concentrations. Indeed, 2.5 μM hemicomplexes assemble to near completion in only 10 minutes. We chose to see if the presence of CCDC32 incubated with 0.5 μM hemicomplexes altered the assembly reaction. Surprisingly, we found that a 1:1 molar equivalent of CCDC32 was a potent assembly *inhibitor* in a time-course assembly assay, with less than 1/3 the total AP-2 assembled in a 60-minute reaction compared to assembly in the absence of CCDC32 (Fig. 4B). Using an end-point assembly assay with varying concentrations of CCDC32, we find that inhibition is concentration dependent, with 50 nm CCDC32 (1/10 molar equivalent) having no effect (Fig. 4C). Inhibition is specific to interaction with the α/σ2 hemicomplex, as a mutant lacking AH1 does not inhibit the assembly reaction (Fig. 4C). Overall, our data, contrary to cellular and genetic experiments, suggest that CCDC32 is a potent inhibitor of AP-2 assembly in solution.

### CCDC32 isolates the α/σ2 from assembled AP-2 in vitro

As CCDC32 is proposed to be an AP-2 assembly factor, it would reason that CCDC32 would have low affinity for assembled AP-2 tetramers (i.e. the product of the assembly reaction) compared to the α/σ2 hemicomplex (i.e. the reactant of the assembly reaction). To test this, we performed Hal-tag pull down assays with CCDC32 and pre-formed AP-2 heterotetramers. Unexpectedly, we found that CCDC32 robustly binds to AP-2 complexes (Figure 4D). This appears to be a conserved phenomenon, as divergent species of CCDC32 were all able to bind to mouse AP-2 core (Supp. Fig. 8C). We next used a panel of mutations and truncations to probe the molecular details of the interaction between CCDC32 and AP-2 core. Interestingly, while the dileucine motif was not necessary for binding α/σ2, the dileucine motif was necessary for binding AP-2 core, and a truncation of CCDC32 containing the dileucine motif (aa150-179) was sufficient for binding (Figure 4D). Truncations of CCDC32 containing only AH1+2 (aa66-130) bound AP-2 core much more weakly than α/σ2 (Fig. 4E). As all of these binding sites should be occluded in the closed AP-2 complex, it suggests that CCDC32 can stimulate opening of AP-2 *in vitro*, similar to other AP-2 binding proteins^36,40^.

We noticed that the β2 subunit consistently appeared sub-stoichiometric compared to the α subunit in pull-downs with CCDC32, including when different species of CCDC32 were used as bait for *M. musculus* AP-2 (Figure 4D, Supp. Fig. 8C). Given that, in cells, CCDC32 is thought to facilitate the interaction of the two AP-2 hemi-complexes, we suspected that this reflected the ability of CCDC32 to drive the reverse phenomenon *in vitro* through Le Chatelier’s Principle^41^. To test this, we used mass photometry to observe the size of AP-2 molecules alone and when co-incubated with CCDC32. AP-2 core alone had a predominant peak at its predicted molecular weight of 203 kDa (Figure 4F, Supp. Fig. 8E,F). However, when co-incubated, a new peak was observed at a molecular weight consistent with the AP-2 hemi-complexes, indicating CCDC32 can isolate AP-2 α/σ2 hemicomplexes *in vitro* (Figure 4F). Further, this phenomenon scaled with the molar ratio of CCDC32 to AP-2 core (supp. Fig. 8E). To better quantify this phenomenon, we designed a dual-fluorescently labeled AP-2 core, with a GFP tag on α and an mScarlet tag on β2 (Fig. 4G). We used this to quantify the β2/α ratio by monitoring the mScarlet/GFP ratio in the elution after a Halo-tag pull-down, normalized to the β2/α ratio of the input dual-labeled AP-2. This shows that indeed CCDC32 isolates α/σ2 hemicomplexes from preformed AP-2 tetramers (Fig 4H).

Using our two assays, we found that only CCDC32 truncations containing the dileucine motif and AH1/2 had sub-stoichiometric β2 (Figure 4H). Given AH1 binds α and is predicted to occupy a large surface area that would otherwise bind β2/μ2, we hypothesized that competition for this binding site allosterically modulates the complex, leading to CCDC32 displacement and AP-2 assembly *in vivo*, and β2/μ2 displacement and AP-2 disassembly *in vitro*. Consistent with this, deleting or mutating AH1 resulted in a failure to displace β2/μ2 from AP-2, including a Y77I mutation to mimic the sequence of the alpha tail (Figure 4H; Supp. Fig. 8F). Interestingly, while neither sufficient nor necessary for binding AP-2 in our hands, deleting or mutating AH2 resulted in failed β2/μ2 displacement as well, including a mutation of a lysine in AH2 predicted to form a salt bridge with AP-2 σ2 (Figure 3B; 4H; Supp. Fig. 5B). Including CCDC32 with deleted or mutated AH1/2 in mass photometry confirmed the findings of the pull-downs, suggesting AH1/2 to be important for AP-2 disassembly *in vitro* (Supp. Fig. 8F). Overall, these data suggest in solution, CCDC32 has a preference for stabilizing the α/σ2 hemicomplex, preventing full complex assembly and even acting to drive assembled tetramers into the hemicomplex state. This function is dependent on specific interactions between CCDC32 and binding sites that are only available in the unassembled hemicomplex.

### Cryo-EM structures of CCDC32 + AP-2 core reveals the last steps in AP-2 assembly

As CCDC32 can isolate α/σ2 hemicomplexes from assembled AP-2 *in vitro*, we hypothesized that co-incubation with CCDC32 would allow the capture of multiple assembly intermediates. To visualize the molecular details of CCDC32-mediated assembly of AP-2, we incubated AP-2 core with CCDC32 and determined multiple high-resolution, single-particle cryo-EM structures. This analysis revealed three distinct structures: two structures of CCDC32 bound to AP-2 and a closed AP-2 structure (Fig. 5A, Supp. Fig. 9,10, Table 2; note we collected data from two grids differing in the amount of time CCDC32 was incubated with AP-2, and found the grid with longer incubation of components had significantly fewer closed AP-2 particles). When bound to CCDC32, we observe AP-2 in an open conformation, with the μ2-CTD disordered and CCDC32 visible in the dileucine binding pocket, as well as a “cargo-bound” conformation, which mimics the conformation of AP-2 in vesicles^17^ with both cargo binding pockets occupied (Fig. 5A). We were unable to clearly detect either AP-2 hemicomplex, perhaps due to their small size and potentially flexible nature.

**Table 2.** Cryo-EM data collection and refinement statistics.

|  | **Closed AP-2** | **AP-2 + CCDC32**<br>**(dileucine motif bound)** | **AP-2 + CCDC32 (dileucine**<br>**and WxxPhi motif bound)** |
| --- | --- | --- | --- |
| EMPIAR ID | EMPIAR-XXXXX | EMPIAR-XXXXX | EMPIAR-XXXXX |
| EMDB ID | EMD-71905 | EMD-71906 (Consensus)<br>EMD-71907 (Focus map)<br>EMD-71909 (Combined) | EMD-71910 (Consensus)<br>EMD-71911 (Focus map 1)<br>EMD-71912 (Focus map 2)<br>EMD-71913 (Focus map 3)<br>EMD-71914 (Combined) |
| PDB ID | 9PWA | 9PWB | 9PWC |
| **Data Collection** |  |  |  |
| Microscope | JEOL CRYO ARM 300 II | JEOL CRYO ARM 300 II | JEOL CRYO ARM 300 II |
| Camera | Gatan K3 (CDS mode) | Gatan K3 (CDS mode) | Gatan K3 (CDS mode) |
| Magnification | 80,000x | 80,000x | 80,000x |
| Voltage (kV) | 300 | 300 | 300 |
| Dose Rate (e-/pixel/s) | 9.7 | 9.7 | 9.7 |
| Exposure Time (s) | 2.67 | 2.67 | 2.67 |
| Number of frames | 60 | 60 | 60 |
| Total electron exposure (e-/Å<sup>2</sup>) | 60 | 60 | 60 |
| Defocus range (µm) | 0.8-1.8 | 0.8-1.8 | 0.8-1.8 |
| Pixel size (Å) | 0.657 | 0.657 | 0.657 |
| Number of micrographs | 31,860 | 68,638 | 68,638 |
| **Reconstruction** |  |  |  |
| Image processing package | cryoSPARC v4.6 | cryoSPARC v4.6 | cryoSPARC v4.6 |
| Final particle images (no.) | 427,544 | 503,371 | 97,343 |
| Symmetry imposed | *C1* | *C1* | *C1* |
| Map resolution (Å) | 2.55 | 2.65 | 3.06 |
| FSC threshold | 0.143 | 0.143 | 0.143 |
| Map resolution range (Å) | 2.4 - 5.5 | 2.3 - 6.1 | 2.7-7.8 |
| Map sharpening B-factor (Å<sup>2</sup>) | -104.9 | -94.6 | -65.2 |
| **Model Refinement** |  |  |  |
| Refinement package | Phenix | Phenix | Phenix |
| Model composition |  |  |  |
| Non-hydrogen atoms | 13707 | 11620 | 13819 |
| Protein residues | 1717 | 1457 | 1734 |
| Ligands | 0 | 0 | 0 |
| B factors (Å<sup>2</sup>); min/max/mean |  |  |  |
| Protein | 14.61/116.84/61.14 | 23.07/148.45/57.12 | 19.16/193.66/73.14 |
| R.m.s. deviations |  |  |  |
| Bond lengths (Å) | 0.003 | 0.005 | 0.004 |
| Bond angles (°) | 0.691 | 0.967 | 0.938 |
| MapCC (volume/mask) | 0.71/0.74 | 0.83/0.82 | 0.82/0.85 |
| **Validation** |  |  |  |
| MolProbity score | 1.68 | 1.54 | 1.57 |
| Clashscore | 6.24 | 7.33 | 7.34 |
| Rotamer outliers (%) | 1.31 | 0.31 | 0.39 |
| Ramachandran Plot |  |  |  |
| Favored (%) | 96.36 | 97.24 | 97.03 |
| Allowed (%) | 3.52 | 2.76 | 2.97 |
| Outliers (%) | 0.12 | 0 | 0 |

The AP-2/CCDC32 open structure shows clear density for the dileucine motif in the dileucine binding pocket of AP-2 α/σ2, consistent with our pull-down data (Fig. 5A,B). This structure is consistent with canonical dileucine cargo engaging an open membrane-bound conformation of AP-2. However, we do not observe a fixed position for the μ2-CTD. Like other dileucine motifs, including CD4 and NEF, CCDC32 makes a salt bridge with R21 of the α subunit, while the LI motif is buried in a hydrophobic pocket containing σ2 residues A63, V88, and V98 (Fig. 5B). CCDC32 makes a second salt bridge with σ2 R15, using an additional acidic residue directly preceding the dileucine motif (Fig. 5B, right). Using fluorescence polarization between AP-2 and Halo-tag CCDC32^150-179^, which contains the dileucine motif, we find a dissociation constant of 0.61 μM (Supp. Fig. 11 A), similar to the ∼0.8 μM affinity of AP-2 for liposomes containing the dileucine motif of CD4^16^. Importantly, we do not observe density for CCDC32 AH1 or AH2 on AP-2. Whereas AH2 would sterically clash with the longin domain of μ2, AH1 should theoretically be compatible with AP-2 bowl binding. Instead, we see the alpha tail packed against α. Therefore, in the open conformation, the only CCDC32-AP-2 interaction is mediated by the dileucine cargo binding motif.

In our previous work, we showed that AP-2 structures do not show density for the μ2-CTD unless tyrosine cargo is included^40,42^. Interestingly, focused 3D classification of our open AP-2/CCDC32 structure showed a population of particles with density consistent with the μ2 CTD (Fig. 5A, middle). This structure closely matches the conformation of AP-2 that is observed in assembled AP-2/clathrin lattices^17^. Given CCDC32 binds the μ2-CTD, we hypothesized this represented a population of molecules with stabilizing interactions between CCDC32 and the μ2-CTD. Indeed, in this structure, we identified additional density occupying the PIP2 and tyrosine cargo binding sites of μ2-CTD (Fig. 5A). Interestingly, CCDC32 contains a WAPL motif that conforms to a Wxxϕ sequence that has been previously reported to bind to the μ2-CTD tyrosine cargo pocket^36,43^. Using AlphaFold3 to model CCDC32 and the μ2-CTD, the only high-confidence interaction shows the WAPL motif binding to the cargo binding pocket (Fig. 5C). Comparison of the Yxxϕ motif of Tgn38 and the Wxxϕ motif of Fcho2 shows that CCDC32 uses a nearly identical mode of binding (Fig. 5C). Furthermore, we observe low-resolution density at both the α and μ2 PIP2 binding pockets, demonstrating the multivalency that underpins CCDC32 binding to AP-2 (Fig. 5A). As with the open structure, we do not observe density for AH1 or AH2 in the cargo-bound conformation, demonstrating the likely stepwise interaction of CCDC32 throughout the assembly process. This matches an AlphaFold prediction with AP-2 core and CCDC32 showing only high-confidence interaction at the two cargo binding sites with the dileucine and WAPL motifs, but no interaction with AH1 or AH2 (Supp. Fig. 6D). Overall, our structural analysis shows that CCDC32 binds to canonical cargo and membrane binding sites to drive AP-2 in the cargo-bound conformation observed on membranes.

### Membrane binding releases inhibitory interactions between CCDC32 and α/σ2 and drives full AP-2 complex assembly

Given our data that IP6 can disrupt interaction of AP-2 μ2 and CCDC32, and we observe low-resolution density at the α and μ2 PIP2 binding pockets, we hypothesized that this may be part of a mechanism where AP-2 sequentially binds CCDC32 and then the plasma membrane during assembly. While CCDC32 has been reported to be both cytosolic^9^ and at clathrin structures at the membrane^31^, it remains unclear if CCDC32 may also have an intrinsic ability to bind membranes. The presence of two amphipathic helices, which are common membrane-binding motifs^44^, supports the idea that CCDC32 directly binds membranes. To test this, we used supported lipid bilayers (SLBs) to perform membrane-binding assays, as we have previously used with other endocytic proteins that bind to AP-2^40^. A schematic of the assay is shown in Supp. Fig. 11B. When incubated with PIP2-containing membranes, Halo-tagged CCDC32 is robustly recruited to the membrane (Fig. 6A, Supp. Fig. 11C-E), while a Halo-tag only control shows no interaction (Fig. 6B, Supp. Fig. 11F). PIP2 may impact CCDC32 binding, as binding is reduced on membranes lacking PIP2 (Supp. Fig. 11C,D). Pelleting assays show that CCDC32 can be washed away from membranes with a buffer containing 0.1% Triton X-100, with no effect when washed in a high salt buffer (Supp. Fig. 11G). As a control, the same assay performed with the BAR domain of Fcho2, which largely relies on electrostatic interactions, is washed away by high salt but unaffected by detergent-containing buffer (Supp. Fig. 11H). Furthermore, an aa150-179 truncation of CCDC2, which does not contain AH1 or AH2, shows no binding (Supp. Fig. 11E). Overall, these data suggest that CCDC32 directly interacts with PIP2-containing membranes using one or both of its amphipathic helices, likely through an insertional mechanism into the outer leaflet.

As AP-2 and CCDC32 both can interact with membranes, we reasoned that the presence of a membrane support might alter our biochemical assays. To test the ability of membranes containing PIP2 to complete AP-2 assembly, we performed Halo-tagged CCDC32 pulldowns of AP-2 in the presence of membranes containing either phosphatidylcholine (PC) and phosphatidylserine (PS), or PC/PS supplemented with PIP2 (Figure 6 C,D). In the presence of PC/PS, the results largely mimic the results with soluble components, where CCDC32 isolates the α/σ2 hemicomplex, with binding broken by an LI->AA mutant, and hemicomplex isolation broken by an AH1 mutant (Figure 6C,D). In the presence of PIP2, however, binding between CCDC32 and AP-2 is strongly stimulated, with all conditions having a roughly stoichiometric amount of all AP-2 subunits. To better quantify AP-2 assembly, we performed the same pulldowns with dual-labeled fluorescent AP-2 and measured the ratio of β2/α (Figure 6E,F). Four of the 5 CCDC32 orthologs show the same behavior when pulling down mouse AP-2 in the presence of membranes (Supp. Fig. 11J). This shows that the presence of PIP2 membranes can strongly drive the reaction to the fully assembled AP-2 state, in contrast to AAGAB which still shows a preference for α/σ2 hemicomplexes (Fig 6F). As AH1 and AH2 both bind to sites on α/σ2 that would preclude full complex assembly, a model emerges whereby CCDC32 holds α/σ2 in an inactive conformation until membrane binding of the AH1 and AH2 domains releases sites for full complex assembly. Membrane-bound α/σ2/CCDC32 ternary complex would not be able to proceed with endocytosis until β2/μ2 recruitment. This model suggests that final assembly of CCDC32-bound AP-2 is driven to completion through interaction with the plasma membrane.

## Discussion

Using a combination of *in vitro* reconstitution and integrative structural modeling, we define the molecular determinants of CCDC32-mediated assembly of the AP-2 clathrin adaptor complex (Fig. 7). In this model, CCDC32 first interacts with AP-2 via strong interactions at the α appendage domain using a novel extended FxDxF motif. Competition for the dileucine cargo binding site on AP-2 leads to AAGAB displacement and stable association of CCDC32 with the α/σ2 hemicomplex. Based on its inhibitory nature in AP-2 assembly assays, we propose that the α/σ2/CCDC32 ternary complex is prevented from full assembly, largely based on competition for binding sites on α via the two amphipathic helices in CCDC32. Assembly inhibition is thereby released upon membrane recruitment, where the AH domains of CCDC32 release from α/σ2 and insert into the membrane. Upon membrane engagement, the intrinsically disordered regions of CCDC32 bind to the μ2-CTD in a multivalent fashion, interacting with both the PIP2 binding site and the tyrosine cargo binding pocket. Binding to the μ2-CTD mediates recruitment of the β2/μ2 hemi-complex, resulting in complex assembly. The final assembly step shows AP-2 in the cargo-bound conformation observed in vesicles, with CCDC32 occupying both cargo binding sites and the PIP2 binding sites on α and μ2. Therefore, co-opting of the most important interaction sites on AP-2 is key to the function of CCDC32.

How CCDC32 is displaced from the AP-2 core is an open question. Closing of the core would, by definition, displace CCDC32, as all CCDC32 binding sites outside of the extended FxDxF motif are occluded in the closed conformation. In this model, assembled AP-2 is released from CCDC32 in the cytosol and then is recruited to endocytic sites via traditional activation pathways. Our data showing that PIP2-containing membranes stabilize assembled AP-2 in the presence of CCDC32 suggest that the plasma membrane may act as the final template for complex assembly, allowing for deposition of AP-2 onto the membrane in the active, cargo-engaged conformation. *In vitro*, absent the membrane, we find that the equilibrium of the assembly reaction does not favor full complex assembly (Fig. 4). The observation that the reaction is driven towards full complex assembly by PIP2-containing membranes strongly suggests that the final stages of AP-2 assembly are mediated by the plasma membrane itself (Figs. 6,7). Alternatively, as AH1 and AH2 use their hydrophobic faces to bind to α/σ2 (Fig. 3B), and these same motifs likely mediate binding of CCDC32 to membranes (Fig. 6A,B; Supp. Fig. 11 B-H), it reasons that membrane localization might trigger CCDC32 displacement, which then allows for β2/μ2 recruitment and full complex assembly. In this way, α/σ2 assembly via AAGAB might be completely cytosolic, while CCDC32 might mediate the final stages of assembly at the membrane. Although CCDC32 was not found to co-localize with AP-2 at mature endocytic sites at the plasma membrane in fixed cells using structured illumination microscopy (SIM)^9^, it was shown to colocalize with clathrin using TIRF^31^. It is possible that the final stages of assembly are transient and only a fraction of AP-2 complexes will be in the assembly process at any given time. Further experiments are required to delineate the temporal ordering of assembly and membrane recruitment in cells, although our results raise the intriguing possibility that the plasma membrane serves as a scaffold for completing the AP-2 assembly reaction.

The presence of a novel extended FxDxF motif in CCDC32 raises the important question of how and when AP-2 is released from this high-affinity interaction. Our data show a binding constant that is at least an order of magnitude higher than any known α appendage binding protein, which typically have dissociation constants in the range of 10-100 μM^37,45^. It stands to reason that competition with other endocytic proteins with an FxDxF domain would be needed to displace CCDC32 from the α appendage, likely at endocytic sites with high concentrations of endocytic regulators. Some proteins, like AP180, have multiple appendage binding motifs, suggesting that multivalency of low-affinity motifs is a hallmark of endocytic proteins^46,47^. However, interaction of these endocytic factors with AP-2 are thought to occur at the plasma membrane, and AAGAB and CCDC32 are proposed to be cytosolic co-factors of AP-2.

This may further support a role for the plasma membrane in AP-2 assembly, if at least to displace CCDC32’s strong interaction with the AP-2 α appendage. Moreover, all of the *in vitro* studies of AP-2 with AAGAB and CCDC32 have utilized an AP-2 construct lacking both appendages. While our data show that the same molecule of CCDC32 is able to bind to α/σ2 and the α appendage simultaneously (Supp. Fig. 4E,F), it is unknown whether there is any allosteric crosstalk between the core and appendage. Further studies with full-length AP-2 subunits will be required to fully explore this possibility.

One striking aspect of our data is that at equimolar concentrations, CCDC32 is able to isolate α/σ2 hemicomplexes from assembled AP-2. In other words, CCDC32 can act as a disassembly factor under certain conditions, which represents the reverse reaction of its role as an assembly chaperone. While this may be the result of the specific experimental conditions used in this study, which lack the same conditions of a crowded cytoplasm, it is possible that disassembly of AP-2 represents another function of CCDC32 in cells. For example, transient disassembly and re-assembly may underpin AP-2 recycling. Importantly, a non-overlapping function between the two AP-2 hemi-complexes has been reported during synaptic vesicle recycling^48^, which would be aided by CCDC32 if it can act as a disassembly factor in certain contexts. Assembly and disassembly is important for regulation of many complexes, including clathrin^49^, which may be the case for the AP-2 complex itself.

While the dileucine binding site was previously known to interact with AAGAB^8^, it is striking that CCDC32 interacts with the majority of known endocytic interaction sites on AP-2. Our structural data show that CCDC32 directly binds to the FxDxF binding site, the dileucine cargo binding site, the tyrosine cargo binding site, and the two highest affinity PIP2 binding sites. From the viewpoint of CCDC32 as an assembly chaperone, this suggests that assembly co-opts known interaction sites, thereby preventing AP-2 from interacting aberrantly with cargo during assembly. This may be a general function of assembly chaperones, whereby the same factor could act to transiently disable subunits from acting outside the context of the assembled complex, while simultaneously templating complex formation. On the other hand, binding to important interaction sites would allow assembly chaperones to have a dual function as more traditional regulatory co-factors. Indeed, CCDC32 has been proposed to be a general endocytic regulator^31^. Future work will be required to delineate the role of CCDC32 as a potential dual-function assembly chaperone and regulator, and to determine if the mechanisms employed in assembling cargo adaptor complexes are applicable to other multi-subunit complexes that require assembly chaperones.

Finally, the cellular roles of AAGAB and CCDC32 as AP-2 assembly chaperones are inferred primarily by biochemical reconstitution assays and reduced AP-2 subunit concentrations in cultured cell lines. The physiological consequences of loss of AAGAB, CCDC32, and AP-2, however, are non-overlapping. Deletion of the *AP2B1* gene in mice causes failed embryo development with cleft palate^50^. Mutation of CCDC32 does not prevent embryonic development, but results in patients with craniofacial malformations^28–30^. Mutation of AAGAB, on the other hand, results in a skin disease with apparently normal craniofacial features^24,25^. This may suggest that the cellular roles of AAGAB and CCDC32 may be nuanced, being either specific to certain cell types, generally less critical for regulating AP-2 compared to cultured cells, or requiring additional co-factors to aid in AP-2 assembly. Future studies in animal models will be necessary to delineate these possibilities.

## Acknowledgements

We thank the lab of Dr. Gunther Hollopeter (Cornell University) for providing plasmids and reagents and the lab of Dr. Amy Gladfelter (Duke University) for use of equipment. We thank Drs. Frederick Hughon, Gunther Hollopeter, Morgan DeSantis, and Michael Cianfrocco for critical reading of the manuscript. Crystallography was performed in the University of North Carolina at Chapel Hill (UNC-CH) Protein Expression and Purification & Macromolecular Crystallography core facility. Diffraction data were collected in the UNC-CH Department of Chemistry X-ray Core Lab. Cryo-EM experiments were performed in at the University of Tokyo EM Facility, which is supported by the Platform Project for Supporting Drug Discovery and Life Science Research (Basis for Supporting Innovative Drug Discovery and Life Science Research (BINDS)) from the Japan Agency for Medical Research and Development (AMED) under Grant Number JP24ama121002 to M.K. DES was supported through a Japan Society for the Promotion of Science (JSPS) international fellowship. R.B.B. acknowledges institutional support from the Department of Biochemistry and Biophysics and the Dean’s Office at the UNC School of Medicine as well as the Lineberger Comprehensive Cancer Center. J.M.S. acknowledges support from the Molecular and Cellular Biophysics Training Grant (T32GM148376-01A1). We thank Stuart Parnham for assistance with NMR experiments. The NMR data presented in this study were collected in the UNC Biomolecular NMR Laboratory, which receives funding from the National Cancer Institute of the National Institutes of Health under award number P30CA016086. We acknowledge pilot funding from the UNC Core Facilities Advocacy Committee and Office of Research Technologies. The content is solely the responsibility of the authors and does not necessarily represent the official views of the National Institutes of Health.

## Materials and Methods

### Reagents

Protease inhibitor cocktail was made in-house from individual components purchased from Gold Biotechnology, Inc. Affinity resins were purchased from Gold Biotechnology, Inc. and Promega. HRV 3C protease, SUMO protease, and Benzonase were purified in house as described^52^. Lipids were purchased from Avanti Polar Lipids. CCDC32 peptides for crystallization were synthesized by SynPeptide Co, Ltd.

### Protein Expression Vectors

pHalo-HRV (control)

pHalo-HRV-AAGAB

pHalo-HRV-CCDC32 WT

pHalo-HRV-CCDC32 mutAH1 (L78A, L81A, K83A, K84A)

pHalo-HRV-CCDC32 mutAH2 (V95S, M100S, L101S, L104S)

pHalo-HRV-CCDC32 mutAH1+2

pHalo-HRV-CCDC32 K108L

pHalo-HRV-CCDC32 dileucine mutant (L153A, I154A)

pHalo-HRV-CCDC32 mutAH1 + dileucine mutant

pHalo-HRV-CCDC32 W19A

pHalo-HRV-CCDC32 FxDxF mutant (F39A, D41A)

pHalo-HRV-CCDC32 extended FxDxF mutant (W19A, F39A, D41A)

pHalo-HRV-CCDC32 aa 66-93

pHalo-HRV-CCDC32 aa 94-130

pHalo-HRV-CCDC32 aa 66-130

pHalo-HRV-CCDC32 aa 1-150

pHalo-HRV-CCDC32 aa 1-30

pHalo-HRV-CCDC32 aa 31-60

pHalo-HRV-CCDC32 aa 61-90

pHalo-HRV-CCDC32 aa 91-120

pHalo-HRV-CCDC32 aa 121-150

pHalo-HRV-CCDC32 aa 150-179

pHalo-HRV-CCDC32 aa 66-130 mutAH1 (L78A/L81A/K83A/K84)

pHalo-HRV-CCDC32 aa 66-130 mutAH2 (V95A/M100A/L101A/L104A)

pHalo-HRV-CCDC32 aa 66-130 mutAH1+2

pHalo-HRV-CCDC32 aa 66-130 Y77I

pHalo-HRV-CCDC32 WAPL mutant (W68A, L71A)

pHalo-HRV-ceCCDC32 (*Caenorhabditis elegans* CCDC32)

pHalo-HRV-dmCCDC32 (*Drosophila melanogaster* CCDC32)

pHalo-HRV-drCCDC32 (*Danio rerio* CCDC32)

pHalo-HRV-hsCCDC32 (*Homo sapiens* CCDC32) p6xHis-CCDC32

p6xHis-CCDC32 dileucine mutant (L153A, I154A) p6xHis-AAGAB

p6xHis-AAGAB aa 1-177 pGH504 (AP-2 alpha core/sigma)

pGB419 (AP-2 beta core/mu)

pEP303 (AP-2 beta core/mu NTD (aa 1-141))

pAlpha(core)-eGFP-6xHis/sigma

pBeta(core)-mScarlet/mu pBeta-6xHis/mu

MZB006 (SUMO-AP-2 alpha appendage (aa 696-938))

MZB066 (AP-2 beta appendage (aa 705-937)-HRV-GST)

pAP-2 Mu CTD aa122-435

pAP-2 Mu CTD PIP2 binding mutant (K341E, K343E, K345E)

### Bioinformatic database searches

Orthologs of *AP2S1, AAGAB*, and *CCDC32* were identified from NCBI’s nr database using 4 iterations of the PSI-BLAST algorithm with the human amino acid sequence as the initial query. Results were filtered for E-scores less than 1E-5. Organisms without a RefSeq sequenced, annotated genome were excluded from analysis, and the NCBI Tree Viewer web tool was used to generate figures. To analyze sequence features of orthologs, a Matlab script was written to take an input fasta file and search input sequence motifs, outputting fasta files of relevant sequence hits and “visual histograms” noting the frequency of occurrence of a given motif along the length of the protein.

### Recombinant protein purification

#### AP-2 complex purification

AP-2 “core” (α 1-621; β2 residues 1-591; μ2; σ2) and AP-2 “bowl” (α 1-621; β2 residues 1-591; μ2 residues 1-135); σ2) was purified essentially as in ^53^. Briefly, BL21 (DE3) cells were grown in Terrific Broth (TB) to mid-log phase and induced at OD 1.0 with 0.5 mM Isopropyl β-d-1-thiogalactopyranoside (IPTG) at 18°C for 18h. Lysis buffer (50 mM HEPES, pH 8.0, 500 mM NaCl, 2 mM MgCl_2_, 1 mM CaCl_2_, 10% glycerol) was supplemented with 1X protease inhibitor cocktail from a 100X stock immediately before resuspension of the cell pellet. After lysis and clarification, protein was incubated with glutathione resin overnight (Gold Bio), washed extensively with low salt wash buffer (50 mM HEPES, pH 8.0, 500 mM NaCl, 1 mM DTT, 10% glycerol), high salt wash buffer (50 mM HEPES, pH 8.0, 1 M NaCl, 1 mM DTT, 10% glycerol) and eluted overnight with gentle rocking in low salt wash buffer supplemented with GST-HRV protease. Eluted protein was concentrated by centrifugal filtration to 2-5 mg/mL before snap-freezing in liquid nitrogen and storage at -80°C.

#### AP-2 α/σ2 purification

AP-2 α/σ2 was purified similarly to AP-2 core and bowl complexes. The differences are as follows: BL21 cells were grown in Lysogeny Broth (LB) rather than TB, and lysis buffer excluded 2 mM MgCl_2_ and 1 mM CaCl_2_ and included 1 mM PMSF and benzonase. After incubation with glutathione resin, only high salt wash buffer without DTT was applied, and eluted with 50 mM HEPES pH 8.0, 500 mM NaCl, 10% glycerol, 10 mM glutathione. GST-HRV protease was then applied during dialysis in 50 mM HEPES, pH 8.0, 500 mM NaCl, 10% glycerol, and the dialysate was re-applied to glutathione resin to remove residual GST and GST-HRV. Eluted protein was concentrated by centrifugal filtration to 2-5 mg/mL before snap-freezing in liquid nitrogen and storage at -80°C.

#### Halo-Tagged α/σ2 purification

BL21 (DE3) cells transformed with their respective protein expression vectors were grown in LB to mid-log phase and induced at roughly OD ∼0.6 with 1 mM IPTG at 18°C for 18-24h. Cells were pelleted by centrifugation and lysis buffer (50 mM HEPES, pH 8.0, 300 mM NaCl, 5 mM Imidazole, 1x protease inhibitor, 1 mM PMSF) was added at ∼30 mL/L of expression culture. After lysis and centrifugation, lysate was applied to 1-2 mL of nickel resin per liter of cell culture. Resin was washed with base buffer (50 mM Tris, pH 8.0, 500 mM NaCl, 5 mM Imidazole), followed by high-salt buffer (50 mM Tris, pH 8.0, 1 M NaCl, 5 mM Imidazole), followed by elution buffer (50 mM Tris, pH 8.0, 500 mM NaCl, 300 mM Imidazole). Protein was further purified using anion exchange chromatography using a 100 mM – 500 mM salt gradient in a base buffer of 50 mM Tris, pH 8.0, 2 mM EDTA, 1 mM DTT. Eluted protein was concentrated by centrifugal filtration to 2-5 mg/mL before snap-freezing in liquid nitrogen and storage at -80°C.

#### AP2 β2/μ2 (His-β2/μ2) purification

BL21 (DE3) cells transformed with their respective protein expression vectors were grown in LB to mid-log phase and induced at roughly OD ∼0.6 with 0.5 mM IPTG at 18°C for 18-24h. Cells were pelleted by centrifugation and lysis buffer (20 mM HEPES, pH 7.6, 300 mM NaCl, 5 mM Imidazole, 1x protease inhibitor, 1 mM PMSF) was added at ∼30 mL/L of expression culture. Resuspended cells were sonicated and ultra-centrifuged to clarify the lysis. 1-2 mL of Ni-NTA resin (GoldBio) equilibrated in deionized water was added to clarified lysate and incubated for at least 30 minutes while rotating at 4°C. Flow-through was collected, and resin was washed with high-salt buffer (20 mM HEPES, pH 7.6, 1 M NaCl, 5 mM imidazole), followed by low-salt buffer (20 mM HEPES, pH 7.6, 100 mM NaCl, 5 mM imidazole), and elution buffer (20 mM HEPES, pH 7.6, 100 mM NaCl, 300 mM imidazole). Protein was dialyzed overnight in 20 mM HEPES, pH 7.6, 100 mM NaCl, 2 mM EDTA, 1 mM DTT. Protein was concentrated, snap-frozen in liquid nitrogen, and stored at -80°C.

#### AP-2 α-appendage purification

An expression construct was made by fusing AP-2 α residues 695-938 (Mus musculus, UniProt P17427) with an N-terminal 10X-His-SUMO tag. BL21 (DE3) cells transformed with this protein expression vector were grown in LB to mid-log phase and induced at roughly OD ∼0.6 with 0.5 mM IPTG at 18°C for 18-24h. Cells were pelleted by centrifugation and lysis buffer (20 mM HEPES, pH 7.6, 200 mM NaCl, 5 mM Imidazole, 1x protease inhibitor, 1 mM PMSF) was added at ∼30 mL/L of expression culture. Resuspended cells were sonicated and centrifuged to clarify the lysis. 1-2 mL of Ni-NTA resin (GoldBio) was added to clarified lysate and incubated for at least 30 minutes while rotating at 4°C. Flow-through was collected, and resin was washed with high-salt buffer (20 mM HEPES, pH 7.6, 1 M NaCl, 5 mM imidazole), followed by low-salt buffer (20 mM HEPES, pH 7.6, 200 mM NaCl, 5 mM imidazole). Protein was eluted using Sumo (CtH) protease and was dialyzed overnight in 20 mM HEPES, pH 7.6, 100 mM NaCl, 1 mM DTT. Protein was further purified using anion exchange chromatography. For fluorescence anisotropy, protein was dialyzed in 100 mM HEPES pH 7.4, 50 mM NaCl, 2 mM DTT as in ^22^. Protein was concentrated, snap-frozen in liquid nitrogen, and stored at -80°C.

#### Halo-Tagged CCDC32 purification

Halo-tagged CCDC32 constructs were all purified in the same manner, which follows the same protocol as for His-β2/μ2. After elution from the nickel resin and dialysis, proteins were further purified to remove excess Halo tag by anion exchange chromatography on a gradient of 100 mM to 400 mM NaCl (buffered with 20 mM HEPES, pH 7.6, 2 mM EDTA, 1 mM DTT), and purified fractions were pooled, concentrated, snap-frozen in liquid nitrogen, and stored at -80°C.

#### Untagged CCDC32 purification

To purify tagless versions of CCDC32 constructs, Halo-tag CCDC32 protein constructs were grown and purified as described in the previous section. In lieu of elution with imidazole, CCDC32 was cleaved off resin by addition of 50 μL of 6-7 mg/mL GST-HRV and co-incubation overnight with gentle agitation in a cold room. Eluted protein was further purified via anion exchange chromatography following the same protocol of the Halo-tag constructs. Purified fractions were pooled, concentrated, snap-frozen in liquid nitrogen, and stored at -80°C.

#### N^15^-labeled CCDC32 purification

BL21 (DE3) cells transformed with their respective protein expression vectors were grown in M9 to mid-log phase and induced at roughly OD ∼0.6 with 1 mM IPTG at 18°C for 18-24h. Cells were pelleted by centrifugation and lysis buffer (20 mM HEPES, pH 7.5, 300 mM NaCl, 5 mM Imidazole, 1x protease inhibitor, 1 mM PMSF and Benzonase was added at ∼30 mL/L of expression culture. After lysis and centrifugation, lysate was applied to 1-2 mL of nickel resin per liter of cell culture. Resin was washed with base buffer (20 mM HEPES, pH 7.5, 300 mM NaCl, 5 mM Imidazole), followed by high-salt buffer (20 mM HEPES, pH 7.5, 1 M NaCl, 5 mM Imidazole), followed by elution buffer (20 mM HEPES, pH 7.5, 300 mM NaCl, 300 mM Imidazole). 6xHis tag was removed from eluted protein by the addition of 1uM HRV protease and simultaneously dialyzed against a buffer containing 20 mM HEPES, pH 7.5, 100mM NaCl, 2 mM EDTA, 1 mM DTT. Protein was further purified using anion exchange chromatography using a 100 mM – 500 mM salt gradient in a base buffer of 20 mM HEPES, pH 7.5, 2 mM EDTA, 1 mM DTT. Eluted protein was dialyzed against a buffer of 20 mM HEPES, pH 6.9, 300mM NaCl, 2 mM EDTA, 1 mM DTT. Protein was then concentrated by centrifugal filtration to 2-5 mg/mL before snap-freezing in liquid nitrogen and storage at -80°C.

#### Dual-fluorescent tagged AP-2 purification

Dual-fluorescent tagged AP-2 was purified essentially as the non-fluorescently tagged version above, except that buffers contained either 5 mM imidazole or 300 mM (elution) and the sample was applied to a Ni+-NTA column. The NaCl concentration of the eluted dual-labeled AP-2 was diluted to 100 mM with 50 mM HEPES pH 7.6, 2 mM EDTA, 1 mM DTT before adding to the FPLC for anion exchange (100-500 mM NaCl gradient with buffer as just described). Purified fractions were pooled, concentrated, snap-frozen in liquid nitrogen, and stored at -80°C.

### *In vitro* pulldowns

All pull-downs were performed with Halo-tagged “Bait” with an HRV cleavage site between the tag and the protein, and untagged “prey”. Proteins were added together at final concentrations of 2 μM bait and 0.5-5 μM prey, depending on the assay, in a final volume of 100 μL base buffer (20 mM HEPES, pH 7.6, 100 mM NaCl, 1 mM DTT, 0.1% Tween-20). 10 μL of magnetic Halolink resin (20% slurry, Promega) was added to 200 μL tubes and washed 5x in base buffer before proteins were added. Tubes were rotated for ∼1 hour at room temperature for 4 hr or overnight at 4°C. Resin was pelleted in a magnetic stand and washed with base buffer three times. 30 μL base buffer containing GST-HRV protease at ∼0.03 mg/mL was used to elute protein from the resin. Input, unbound, and elution fractions were run on SDS-PAGE gels and visualized by Coomassie blue staining.

#### Halo-CCDC32 + α appendage pull down

The pull-down assay for CCDC32 and the α appendage was quantified by gel densitometry in FIJI^54^. Mean and standard error of the mean was calculated and confidence intervals were calculated using a one-way ANOVA in Prism (GraphPad).

#### AAGAB/CCDC32 competition experiment

Halo-HRV-α/σ2 (2 μM) was rotated with 6xHis-HRV-AAGAB construct (1 μM) and 10 μL Magne-HaloTag beads for one hour at room temperature. 6xHis-HRV-CCDC32 construct (20 μM) was added to solution and rotated for another hour at room temperature. Beads were washed 4x before eluting with HRV protease (0.3 μM) in buffer for one hour at room temperature. The buffer for dilutions and washes was 20mM HEPES, 100mM NaCl, 0.1% v/v Tween-20, 1mM DTT

#### Membrane pull-downs

For the pull-down with liposomes, liposomes with 1 mM total lipid content were added with the bait and prey molecules, either PC/PS (80%, 20%) or PC/PS/PIP2 (80%, 15%, 5%).

#### Dual-labeled fluorescent AP-2 pulldowns

Dual-labeled fluorescent AP-2 pulldowns were done as above. Liposomes were added in the experiments in figure 7F with 0.5 mM total lipid content and either PC/PS/Cy5-PE (79.5%, 20%, 0.5%) or PC/PS/PIP2/Cy5-PE (79.5%, 15%, 5%, 0.5%). Samples were added to a clear bottom 384 well plate (Greiner) and imaged on a PheraStar instrument.

### Membrane binding assays

#### Generation of SLBs

5 mM of small unilamellar vesicles (SUVs) of a desired lipid composition were mixed with 440 mm^2^ of 6.5 micron (diameter) silica microspheres in a supported lipid bilayer (SLB) buffer containing 300 mM NaCl, 20 mM HEPES pH 7.4, and 1 mM MgCl_2_. SUVs and microspheres were rotated using an over-end mixer for 1 hour at room temperature. Lipid-coated microspheres were then spun down at 300g (minimal sedimentation velocity) for 30 seconds. The supernatant was removed and the lipid-coated microspheres were resuspending in wash buffer (100 mM NaCl, 20 mM Hepes pH 7.4, 1 mM DTT). Wash steps were repeated three additional times. 2.5 mm^2^ of lipid-coated microspheres were mixed with fluorescently labeled proteins in a custom-built reaction well glued onto a poly-ethylene glycol passivated coverslip. The reaction was allowed to reach steady state (1 hour) before imaging.

#### Imaging and quantification of fluorescent protein binding to SLBs

Protein binding to SLBs was assessed through imaging on a custom-built spinning disc (Yokogowa) confocal microscope equipped with a Ti-82 Nikon Stage, a 100x Plan Apo 1.49 numerical aperture objective and a Zyla sCMOS camera (Andor). Following image acquisition, raw micrographs were imported into Imaris v.8.1.2 (Bitplane, AG). Images were background subtracted using the software’s Gaussian filter for determining background signal. Each bead was identified using the fluorescence from the lipid channel. Sum intensity values from each fluorescent channel (protein and lipid) at each focal plane were obtained and used to calculate a ratio of protein intensity to lipid intensity.

#### Pelleting binding assay

SLBs were generated as described above and mixed with 5 µM of either Halo-CCDC32 or Fcho2 in low adhesion tubes. The reaction was allowed to reach steady-state (1 hour at room temperature). The SLB-protein mixture was then washed using wash buffer (100 mM NaCl, 20 mM HEPES pH 7.4, 1mM DTT). SLB-protein mixture was then resuspended in a buffer containing 1 M NaCl, 20 mM HEPES pH 7.4, and 1mM DTT or 100mM NaCl, 20 mM HEPES pH 7.4, 1 mM DTT, and 0.1% Triton X-100. The mixture was incubated within the respective buffers for 15 minutes. The mixture was then spun down from which supernatant and pellet fractions were obtained for SDS-PAGE analysis.

### Circular Dichroism Spectroscopy

Circular Dichroism spectra were obtained using a Chirascan V100 Spectrometer (Applied Photophysics). Spectra were obtained using 5 µM of CCDC32 in 100 mM NaF, 10 mM Potassium Phosphate pH 7.4, and 1 mM TCEP in a 1-mm pathlength quartz. Measurements were taken from 185 nm through 260 nm.

### Fluorescence Polarization

384-well plates were coated in 0.2 mg/mL BSA overnight at 4°C Celsius. The following day, BSA solution was removed and washed three times using 100 mM NaCl, 20 mM HEPES pH 7.4, 1 mM DTT. 5 nM of Atto488-labeled Halo-CCDC32^150-179^ was mixed with various concentrations of AP-2 core in a final buffer composition of 100 mM NaCl, 20 mM HEPES pH 7.4, 1 mM DTT. The reaction was allowed to reach steady-state (1 hour at room temperature). Fluorescence polarization measurements were obtained using a CLARIOstar plate reader.

### AlphaFold model prediction

AlphaFold modeling was done in three ways: the AlphaFold3 server and default parameters, the CHAI-1 server using default parameters, and a local install of ColabFold running AlphaFold-multimer v3, with recycles set to 12. Sequences for proteins used are included in figure legends. Visualization of AlphaFold models was done using ChimeraX software.

### Mass photometry

All measurements were taken on a Refeyn One Mass Photometer with AcquireMP software and analyzed using DiscoverMP software (all from Refeyn Ltd, Oxford, UK) according to previously established protocols. Briefly, imaging coverslips were bath sonicated in 50% isopropanol for 15 minutes and subsequently in ultra-pure H_2_O for 5 minutes. Coverslips were dried using a stream of filtered air and imaging wells were made by applying adhesive four-well gaskets (Thor Labs). Prior to imaging, 10 μL of reaction buffer (50 mM HEPES pH 7.6, 500 mM NaCl, 10% glycerol) was added to the imaging well to determine the imaging plane using an internal autofocusing system. 10 μL of sample was added to the imaging buffer, resulting in a final volume of 20 μL and a sample concentration between 5-40 nM. Measurements were acquired for 1 minute at a frame rate of 100 fps. For runs containing binding partners, proteins were mixed at a 1:1 molar ratio and allowed to equilibrate for at least 30 minutes at room temperature before data collection.

#### Assembly assays

Mass photometry assembly assays were quantified by determining the relative ratio of assembled to unassembled peak counts (see Figure 4A). Given apparent dimerization of β2/μ2 artificially increasing the assembled peak counts (see Supp. Fig. 8A), ratios were fitted to a linear curve where the mean ratio of 50 nM hemi-complex assembly at 0 min was treated as 0% assembly, and the mean ratio of 2.5 μM hemi-complex assembly at 60 min was treated as 100% assembly.

### NMR spectroscopy

NMR experiments were carried out at 298 K on a Bruker Avance III 850 MHz spectrometer equipped with a cryogenic probe. A 100 µM sample of ^15^N-CCDC32 was prepared in buffer containing 20 mM HEPES pH 6.9, 100 mM NaCl, 2 mM EDTA, 1 mM DTT, and 1% D2O. ^1^H-^15^N HSQC spectra were collected using standard Bruker pulse sequences. NMR data were processed using NMRPipe^55^ and analyzed using NMRFAM-SPARKY^56^ on NMRbox^57^.

### X-ray crystallography structure determination

#### Crystallization and data collection

Purified AP-2 α-appendage was mixed with synthesized CCDC32 peptides for crystallization. For the FxDxF peptide (CCDC32 residues 37-45; sequence: NAFSDSFMD), final protein concentration was 4.8 mg/ml in 20mM HEPES, pH 7.7, 100mM NaCl, 1mM DTT buffer. Final FxDxF peptide concentration was 4.0 mM. For the extended FxDxF peptide (CCDC32 residues 17-44; sequence: DLWAEICSCLPSPAQEDVSDNAFSDSFM), final protein concentration was 3.0 mg/ml in 20mM HEPES pH 7.7, 150mM NaCl, 5mM Imidazole, 3.5% (v/v) DMSO buffer. Final extended FxDxF peptide concentration was 2.0 mM. Crystallization experiments were set up using 100 nL protein to 100 nL reagent drop ratios in commercially available matrix screens. Crystals of the appendage-extended FxDxF were observed following one day of growth in a condition consisting of 100 mM SPG pH 8.0, 25% PEG 1500 (Molecular Dimensions PACT Premiere, A5). Crystals of the appendage-FxDxF complex were grown similarly against a reservoir of 40% isopropanol, 100 mM imidazole pH 6.5, 15% PEG 8000 (Rigaku Reagents Wizard Classic 3 and 4, H9). For cryo-protection, crystals were quickly treated with a soak of 15% ethylene glycol then flash frozen in liquid nitrogen. Diffraction data were collected on a Bruker D8 Venture using an IµS copper (K-α) sealed microfocus source and a PHOTON III C14 detector. Crystals were kept frozen with an Oxford Cryostream 800. Data were reduced in Saint and scaled in SADABS^58^.

#### Data processing and model building

The structures were phased by molecular replacement using Phenix Phaser with PDB 1W80 as a starting model based on sequence homology^37,59^. The structures were treated to alternating rounds of reciprocal space refinement in Phenix and real space rebuilding in Coot^60–63^. PEG polymer ligands were built using eLBOW^60^. The protein chains were numbered following mouse AP2A2 (UniProt P17427) and the peptide chains by mouse CCDC32 (UniProt Q8BS39). The structures were deposited in the Protein Data Bank under accession numbers 9PUE and 9PPP.

### Cryo-EM structure determination

#### Sample preparation

UltrAuFoil R 1.2/1.3 300 mesh grids were washed in acetone and glow-discharged before sample application. AP-2 core at ∼0.4 mg/mL was incubated with 5x molar excess of 6xHis-CCDC32 for ∼30 min before freezing in the following buffer: 20 mM HEPES pH 7.6, 180 mM NaCl, 2% glycerol, 1 mM DTT, 0.07% octyl-beta-glucoside. 2 μL sample was applied to grid and blotted once manually. Sample was reapplied and blotted before freezing in liquid ethane held at −190 °C using a Vitrobot Mark IV (ThermoFisher) with the following settings: 2 μL of sample, 4 °C, 100% humidity, blot force −10 and blot time 4 s.

#### Data collection

Images were recorded on a JEM-3300 (CRYO ARM™ 300 II, JEOL, Tokyo, Japan) at the University of Tokyo, operated at 300 keV. An Omega filter with a slit width of 20 eV and a Gatan K3 direct electron detector in correlated-double sampling (CDS) mode were used for imaging. Beam tilt calibration was performed in SerialEM^64^. The nominal magnification was set to 80,000×, yielding a physical pixel size of 0.657 Å/pixel. Movies were acquired using SerialEM, with a target defocus range of 0.8–1.8 µm. Each movie was recorded for 2.40-2.67 s with a total electron dose of 60 e?/Å^2^, divided into 60 frames. Two datasets were collected on the same microscope and with identical parameters (31,860 micrographs and 36,778).

#### Structure determination

All processing was performed in cryoSPARC v4.6.0. Each dataset was processed independently with the following schema: patch-based motion correction with dose weighting, patch-based CTF estimation, blob picking for initial particle picks, 2D classification, template picking with class averages, 2D classification. “Clean” particle picks were classified using *ab-initio* model generation with multiple classes, followed by heterogeneous refinement with the references generated in the *ab-initio* stage. This yielded two high-resolution classes: an open conformation of AP-2 with density for CCDC32 visible at the dileucine binding pocket, and “closed” AP-2 with no density for CCDC32. After Non-Uniform refinement, a Topaz model was trained for each particle class, and the micrographs were re-picked. *Abinitio* model generation followed by heterogeneous refinement yielded two high-resolution classes, albeit with more particles. For “closed” AP-2, a final dataset of 427,544 particles was refined, polished using reference-based motion correction, followed by a final Non-Uniform refinement, including refinement of defocus and global CTF parameters. This yielded a final reconstruction with a gold-standard Fourier shell correlation (GSFSC) resolution of 2.55 Å. For the “open” AP-2 particles, a second Topaz model was trained, followed by *ab-initio* model generation and heterogeneous refinement, yielding a final class of 503,371 particles. Particles were refined, polished using reference-based motion correction, followed by a final Non-Uniform refinement, including refinement of defocus and global CTF parameters. This yielded a final reconstruction with a GSFSC resolution of 2.65 Å. To improve the resolution of local features of this map, two rounds of local refinement was performed, first using a loose mask around the α/σ2 interface and standard local refinement parameters and second with a tighter mask and more constrained search parameters. This dramatically improved the resolution of this region of the complex. Half maps were combined using phenix.combine_focused_maps (phenix^65^ version 1.21.1-5286), followed by map sharpening of the composite half-maps.

During our processing, we noticed that some 3D classes of AP-2 in the open conformation had additional, low-resolution density. Additional processing to increase the number of this particle class yielded a highly-anisotropic reconstruction of AP-2 in the “cargo-bound” conformation, nearly identical to 2xa7.pdb, a crystal structure of AP-2 bound to the tyrosine-cargo motif of Tgn38. The anisotropy of these particles could not be improved. Instead, we used a 3D mask around the region of presumptive density in our open AP-2 reconstruction to perform focused 3D classification. This analysis showed that ∼15% of our open AP-2 particles were indeed in the cargo-bound conformation, with no anisotropy in the map from directional orientation bias. To increase the number of particles, we re-processed both datasets using blob picker, template picker, and Topaz picker, followed by extensive 2D and 3D classification. Duplicate particles were removed, yielding a combined dataset of ∼1.7 million particles of AP-2 in the open conformation. Focused 3D classification looking for cargo-bound AP-2 particles showed several cargo-bound classes, with a total particle count of 284,383 particles. These particles were refined, showing a GSFSC resolution of 2.91 Å, but with variable resolution, particularly in the AP-2 cargo-binding domain. In lieu of focused refinement, a final round of focused 3D classification revealed a class containing 97,343 particles with a GSFSC resolution of 3.06 Å, but with far better density for the cargo-binding domain. This was further improved using three separate focused refinements. Half maps were combined using phenix.combine_focused_maps, followed by map sharpening of the composite half-maps.

### Model building and validation

#### Closed AP-2

The X-ray structure of closed AP-2 core (2VGL.pdb^12^) was docked into the map and all solvent and co-factors were removed. The 4 protein chains were re-docked using the fit-in-map function in ChimeraX to account for slight structural changes. The model was visually inspected and manually rebuilt where needed in Coot, followed by phenix.real-space_refinement, including B-factor refinement.

#### Open, dileucine-bound AP-2

The primary sequence for AP-2 (α 1-620, β2 1-591, μ2 1-135, σ2) was run as a prediction in AlphaFold2-multimer-v3 with full-length mouse CCDC32 (UniProt Q8BS39). Multiple models showed high-confidence interaction of the dileucine motif (aa 154-159) bound in the correct binding pocket on AP-2. The top model was docked into the cryo-EM map and regions of t model were manually re-docked using the fit-in-map function in Chimera. The model was visually inspected and manually rebuilt where needed in Coot, followed by phenix.real-space_refinement, including B-factor refinement.

#### Cargo-bound AP-2

The primary sequence for AP-2 (α 1-620, β2 1-591, μ2, σ2) was run as a prediction in AlphaFold2-multimer-v3 with full-length mouse CCDC32 (UniProt Q8BS39). In general, models showed AP-2 in the correct conformation, with a high-confidence interaction of the dileucine motif (aa 154-159) bound in the correct binding pocket on AP-2. However, all models showing interaction with the μ2-CTD were low confidence. Modeling of CCDC32 with the μ2-CTD using the AlphaFold3 server showed high-confidence interaction of the “WAPL” motif of CCDC32 (aa68-71) binding to the tyrosine cargo binding pocket on AP-2. The two AlphaFold models were docked into the sharpened, composite map and regions of the model were manually docked using the fit-in-map function in ChimeraX. The model was visually inspected and manually rebuilt where needed in Coot, followed by phenix.real-space_refinement, including B-factor refinement.

---

**Supplementary Figure 1.**
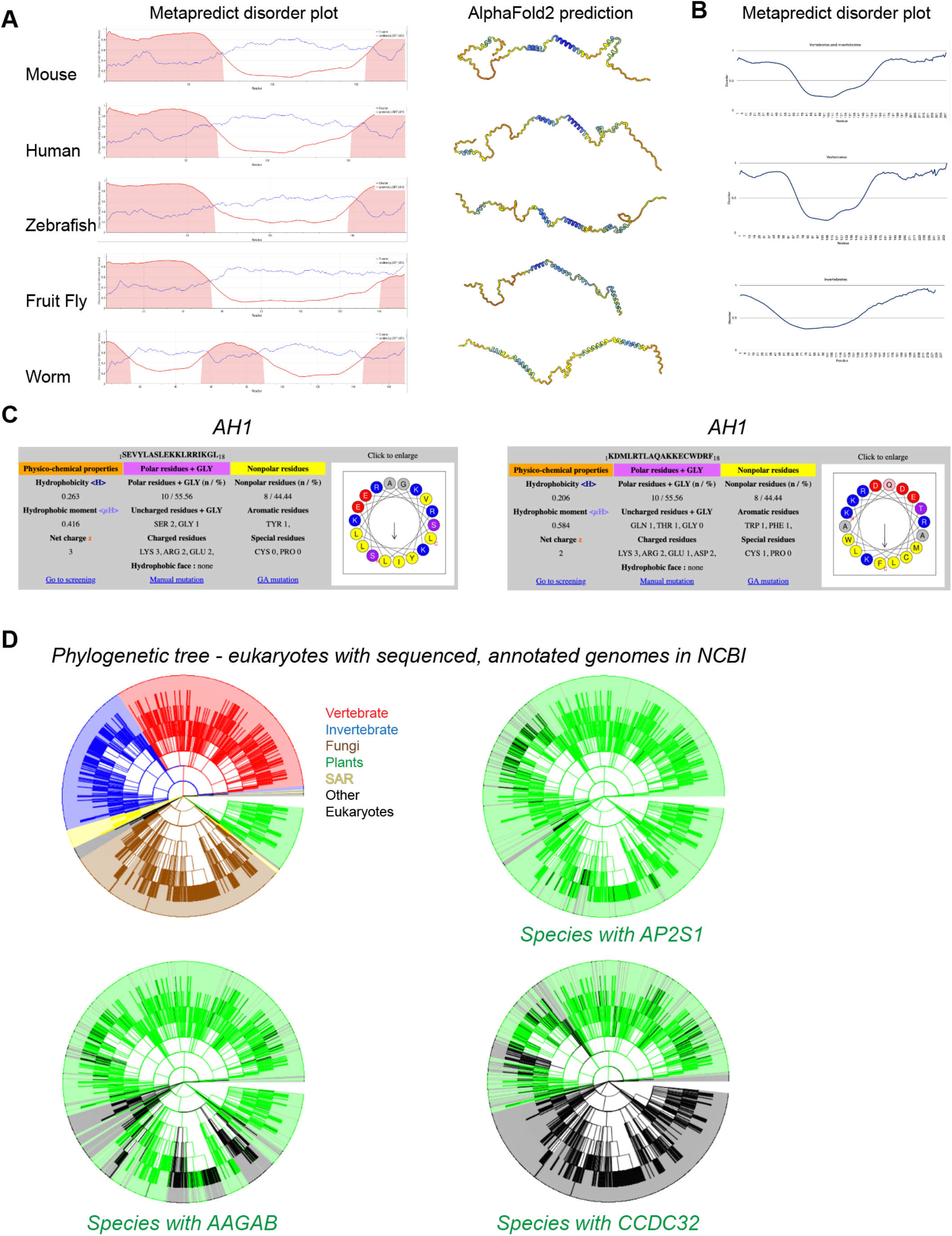
**(A)** A Metapredict disorder plots for five CCDC32 orthologs, showing a prediction of protein disorder based on primary sequence. The Uniprot Alphafold2 prediction for each ortholog is shown on the right, colored by pLDDT score. **(B)** Metapredict disorder plots for all metazoan CCDC32 orthologs, vertebrate orthologs, or non-vertebrate orthologs. Fastas were batch imported into Metapredict and the disorder plot represents an average of all sequences. **(C)** Heliquest plots for CCDC32 AH1 and AH2, showing helical wheel diagrams and calculated hydrophobic moment values. **(D)** Phylogenetic trees showing AP-2, AAGAB, and CCDC32 conservation. First, all sequenced and annotated genomes in NCBI were extracted and built into a phylogenetic tree (rainbow, top left). Then AP-2 σ2, AAGAB, and CCDC32 orthologs were found by sequence homology using PSI-BLAST^51^. Species with high-confidence orthologs were plotted as green.

**Supplementary Figure 2.**
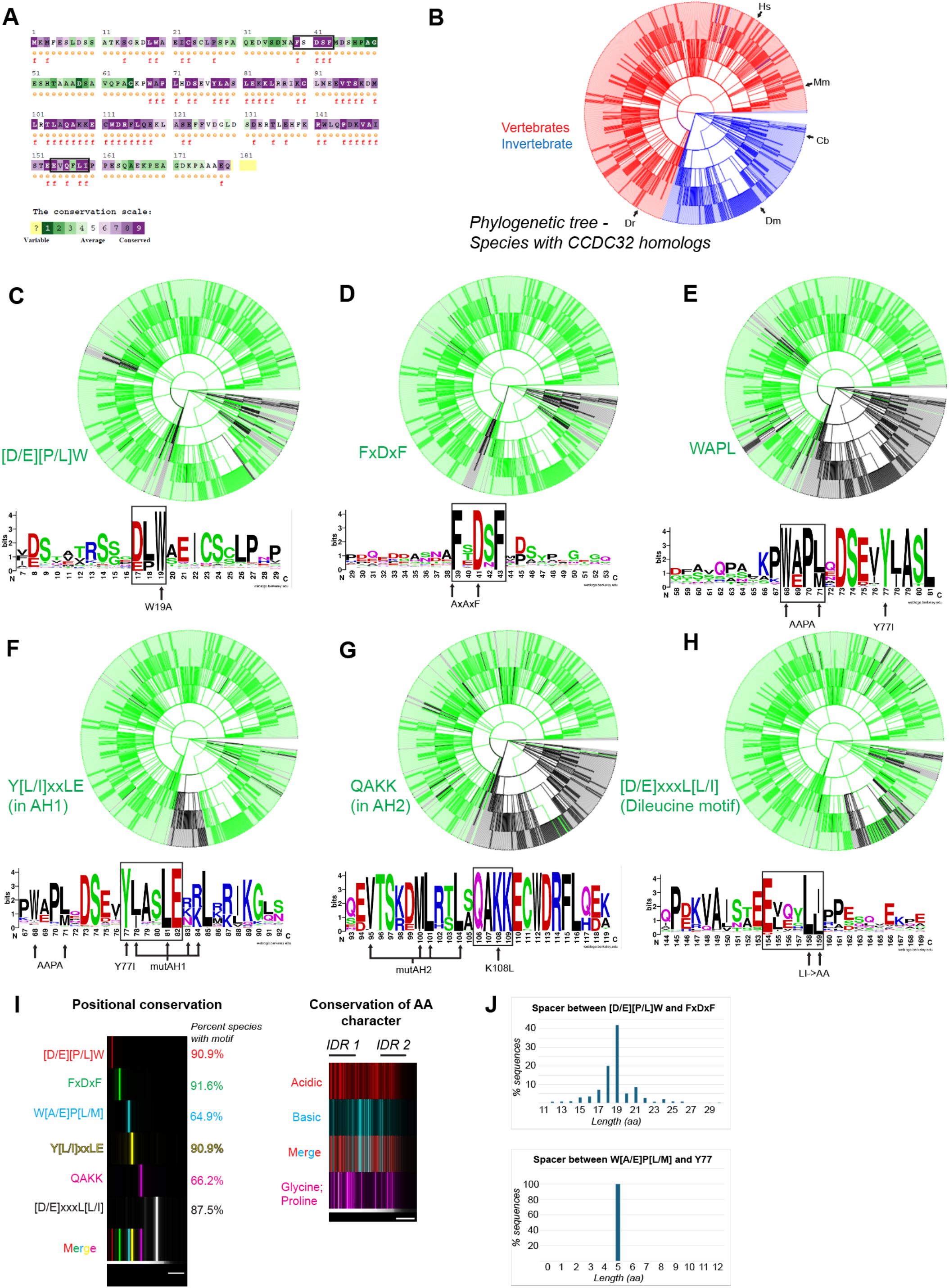
**(A)** Primary sequence conservation of mouse CCDC32, analyzed and colored using the Consurf database. **(B)** Phylogenetic tree of species with CCDC32 orthologs, colored by vertebrates and invertebrates. **(C)** Phylogenetic tree from (**B**) colored green if the ortholog contains a [D/E][P/L]W motif. The amino acid frequency plot centered on this motif is shown below. **(D)** Phylogenetic tree from (**B**) colored green if the ortholog contains an FxDxF motif. The amino acid frequency plot centered on this motif is shown below. **(E)** Phylogenetic tree from (**B**) colored green if the ortholog contains a WAPL motif. The amino acid frequency plot centered on this motif is shown below. **(F)** Phylogenetic tree from (**B**) colored green if the ortholog contains a Y[L/I]xxLE motif, a highly conserved sequence in AH1. The amino acid frequency plot centered on this motif is shown below. Residues mutated to alanine in the mutAH1 construct are labeled. **(G)** Phylogenetic tree from (**B**) colored green if the ortholog contains a QAKK motif, a highly conserved sequence in AH2. The amino acid frequency plot centered on this motif is shown below. Residues mutated to serine in the mutAH2 construct are labeled. **(G)** Phylogenetic tree from (**B**) colored green if the ortholog contains a [D/E]xxxL[L/I] “dileucine” motif. The amino acid frequency plot centered on this motif is shown below. **(H)** Positional sequence conservation plots. All sequences from (**B**) were assigned a black or colored value at each amino acid location according to the motif (Y axis) and then all values were averaged at all amino acid locations. Sequences were not aligned, but rather plotted as raw location values. Left, positional conservation of defined motifs. Right, positional conservation of amino acids by chemical identity. **(I)** Histogram distribution of linker length between sequence motifs. Contains all sequences from (**B**).

**Supplementary Figure 3.**
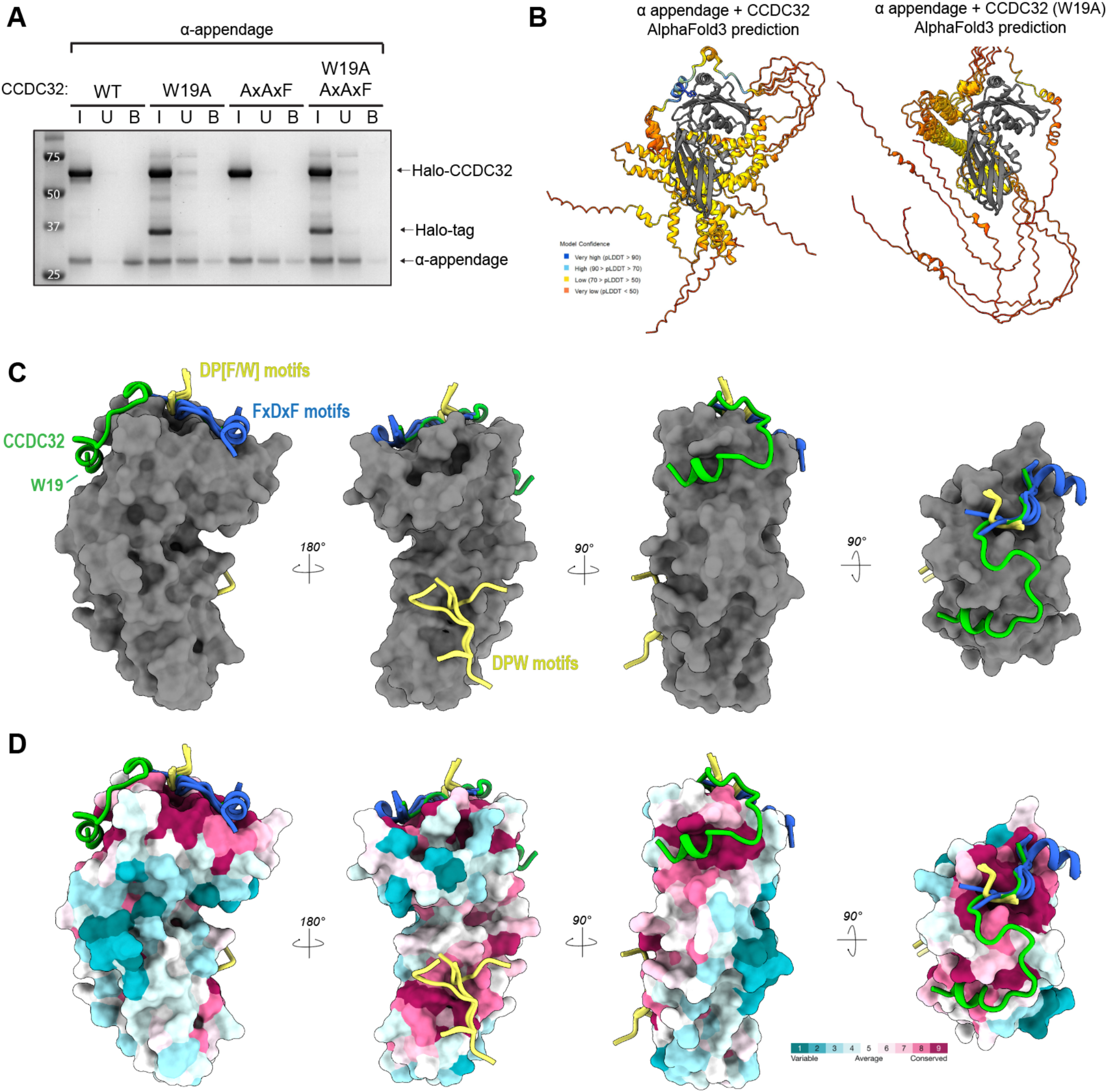
**(A)** Uncropped gel image from the pulldown binding assay between CCDC32 mutants and the α appendage in Figure 2B. Proteins were eluted on the resin by boiling beads, as CCDC32 and α appendage co-migrate on the gel. Halo-Tag CCDC32 remains on the beads as it is covalently linked via the Halo-Tag. **(B)** Alphafold3 predictions of CCDC32 and α appendage, either use WT or W19A sequences. CCDC32 is shown colored by pLDDT score, α is colored grey. **(C)** Alignment of all report α appendage structures with CCDC32-α appendage structure. CCDC32 (green), DPW and DP[F/W] motif peptides (yellow), FxDxF peptides (blue), α (grey). **(D)** Structures as shown in (**C**), but the surface of the α appendage is colored by conservation score, using the Consurf server. Note three discrete conserved surfaces on the α appendage.

**Supplementary Figure 4.**
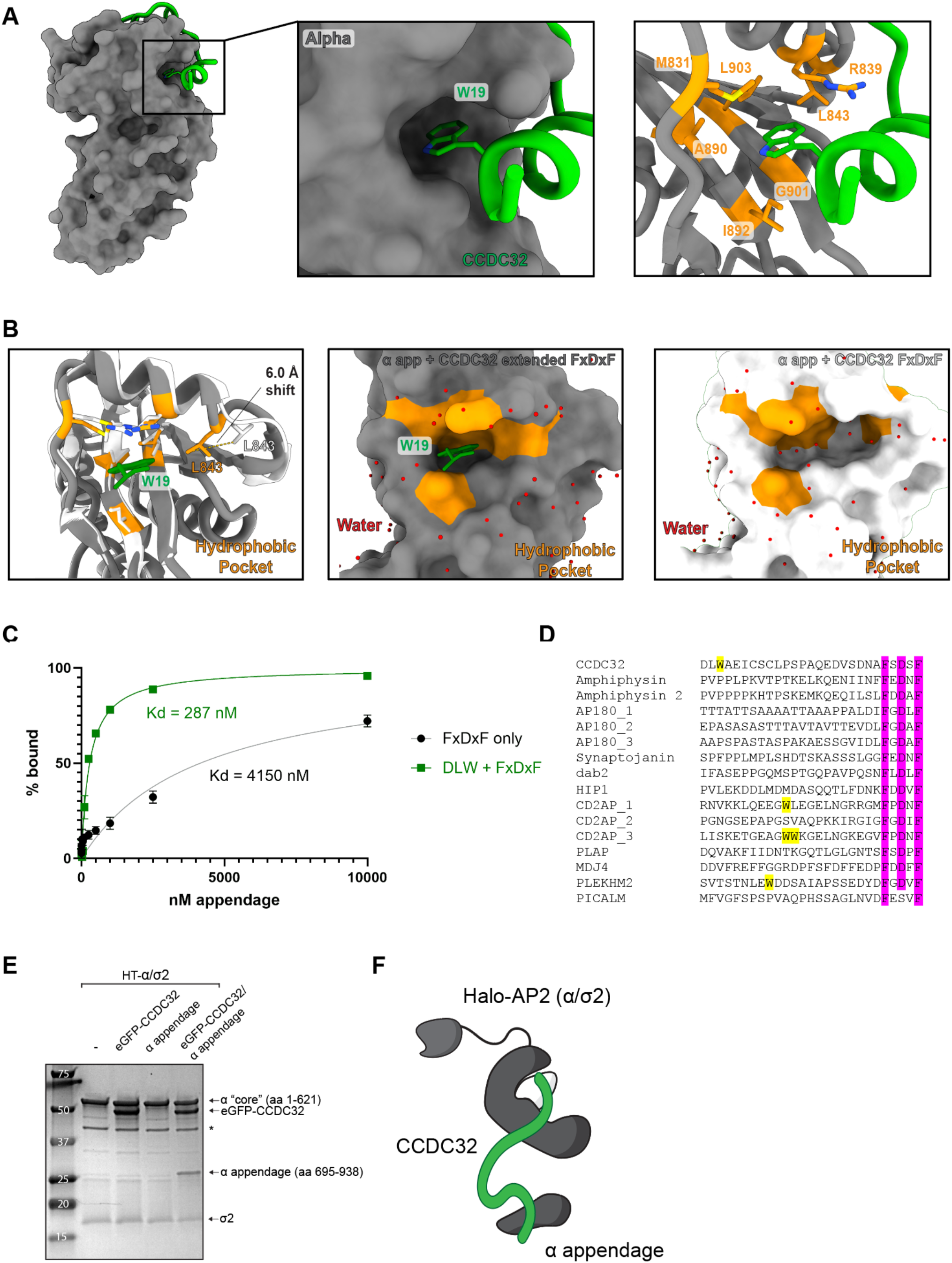
**(A)** Schematic of CCDC32 W19 packing into a hydrophobic pocket on the backside of the α appendage. (Right) hydrophobic residues are shown in stick representation and colored orange. **(B)** Comparison of CCDC32 extended FxDxF crystal structure (grey) and CCDC32 FxDxF crystal structure (white). Not the 6 Å shift of a loop (L843) labeled, which closes a hydrophobic pocket. Compare (middle) with (right), showing that when not occupied by W19, the pocket is much larger and accommodates several water molecules. Binding of W19 closes the pocket and displaces several waters. **(C)** Binding isotherms and fitted Kd values for α appendage binding to FITC-labeled CCDC32 peptides. **(D)** Sequence alignment of reported proteins with FxDxF motif, aligned on the motif. All tryptophans are highlighted in yellow. Sequences come from mouse orthologs. **(E)** Binding assay with HaloTag-α/σ2 bait pulling down eGFP-tagged CCDC32 or α appendage. eGFP-CCDC32 was used as untagged CCDC32 co-migrates on the gel with α appendage. HaloTag-α/σ2 can pull down CCDC32, but not α appendage. In the presence of CCDC32, α appendage is pulled down, showing that one molecule of CCDC32 can bind to the α/σ2 core and the α appendage simultaneously. **(F)** Schematic for pull-down assay showing bridging interaction of CCDC32.

**Supplementary Figure 5.**
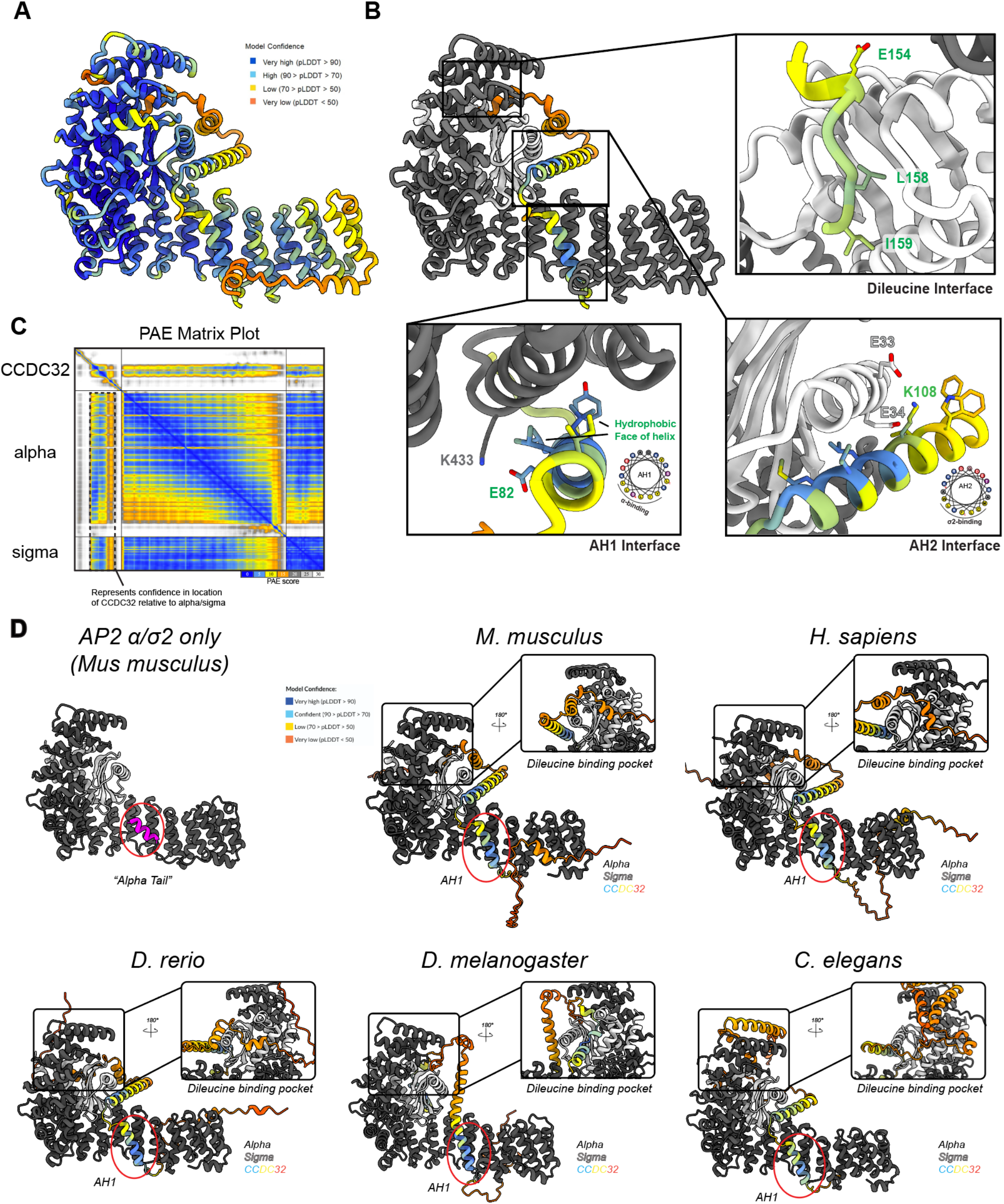
**(A)** AlphaFold3 prediction of the α/σ2 hemicomplex bound to CCDC32. The model is colored according to the pLDDT score. **(B)** AlphaFold3 prediction shown in (**A**), with α/σ2 colored grey/white. Three predicted interfaces (AH1, AH2, dileucine motif) are shown inset. **(C)** Predicted Aligned Error (PAE) matrix plot from the AlphaFold prediction in (**A**). Highlighted in a dashed box is the pairwise confidence between AH1/AH2/dileucine region of CCDC32 and α/σ2. **(D)** AlphaFold3 predictions of α/σ2 from various orthologs. α/σ2 colored grey/white and CCDC32 colored by pLDDT score. An AlphaFold model of α/σ2 is shown in the top left, with the alpha tail colored magenta, showing that it occupies the same binding site as AH1.

**Supplementary Figure 6.**
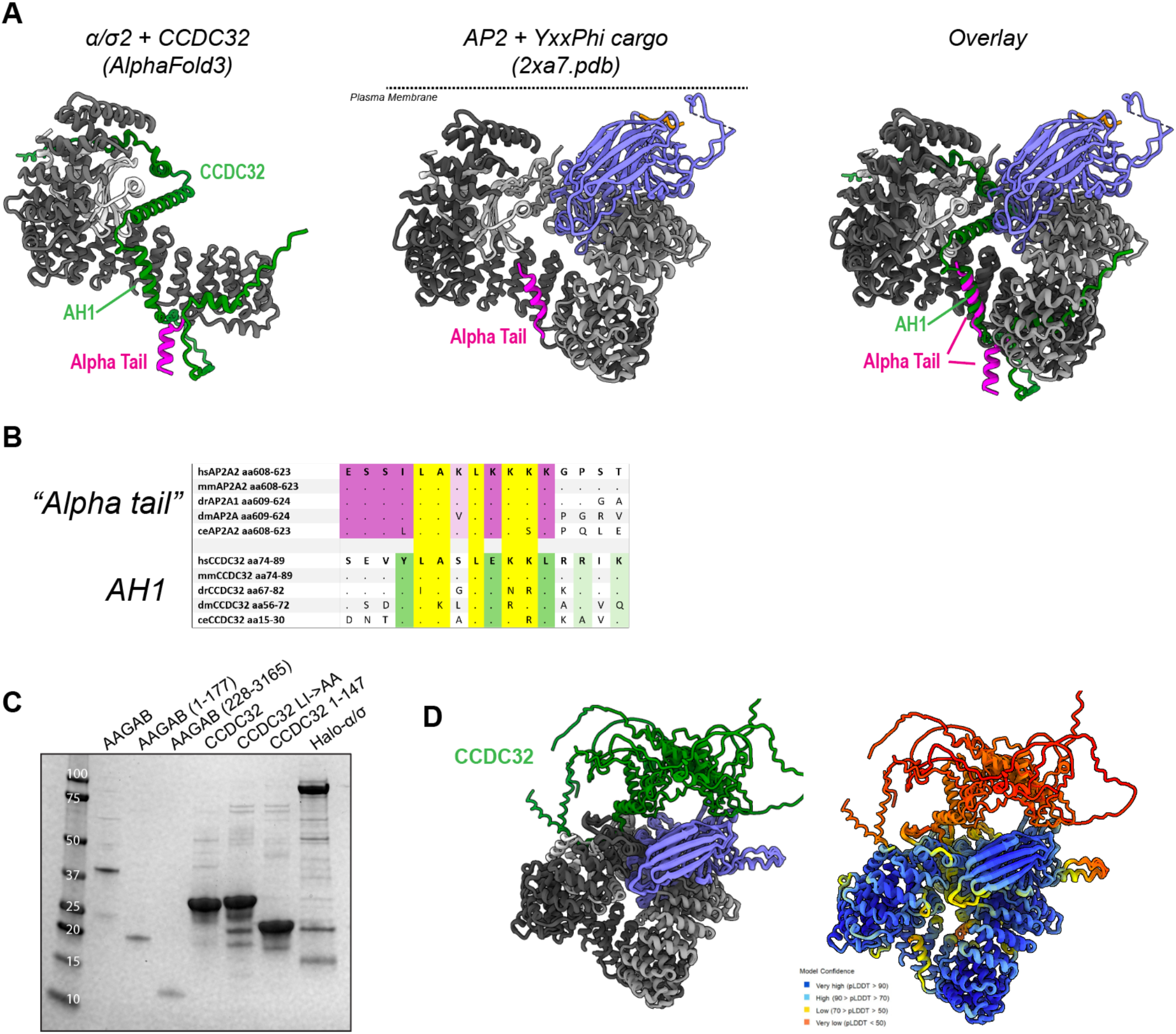
**(A)** Comparison of the AlphaFold3 structure of α/σ2 (left) compared with the crystal structure of AP-2 with Tgn38 tyrosine cargo peptide, 2xa7.pdb (middle). The alpha tail is colored magenta in both. Right, an overlay of the two structures, demonstrating that multiple CCDC32 binding motifs are incompatible with binding assembled AP-2. **(B)** Sequence alignment of the alpha tail from AP-2 α and AH1 From CCDC32. **(C)** SDS-PAGE gel for AAGAB/CCDC32 competition experiment shown in **Fig. 3F**. Note that AAGAB^228-315^ is visible on the input gel, but is not seen in the elution gel, showing that it does not bind to α/σ2 *in vitro*. **(D)** AlphaFold2-multimer v3 prediction of AP-2 core and CCDC32. Note that CCDC32 only contacts the dileucine and tyrosine cargo binding sites and not on α/σ2 via AH1 or AH2.

**Supplementary Figure 7.**
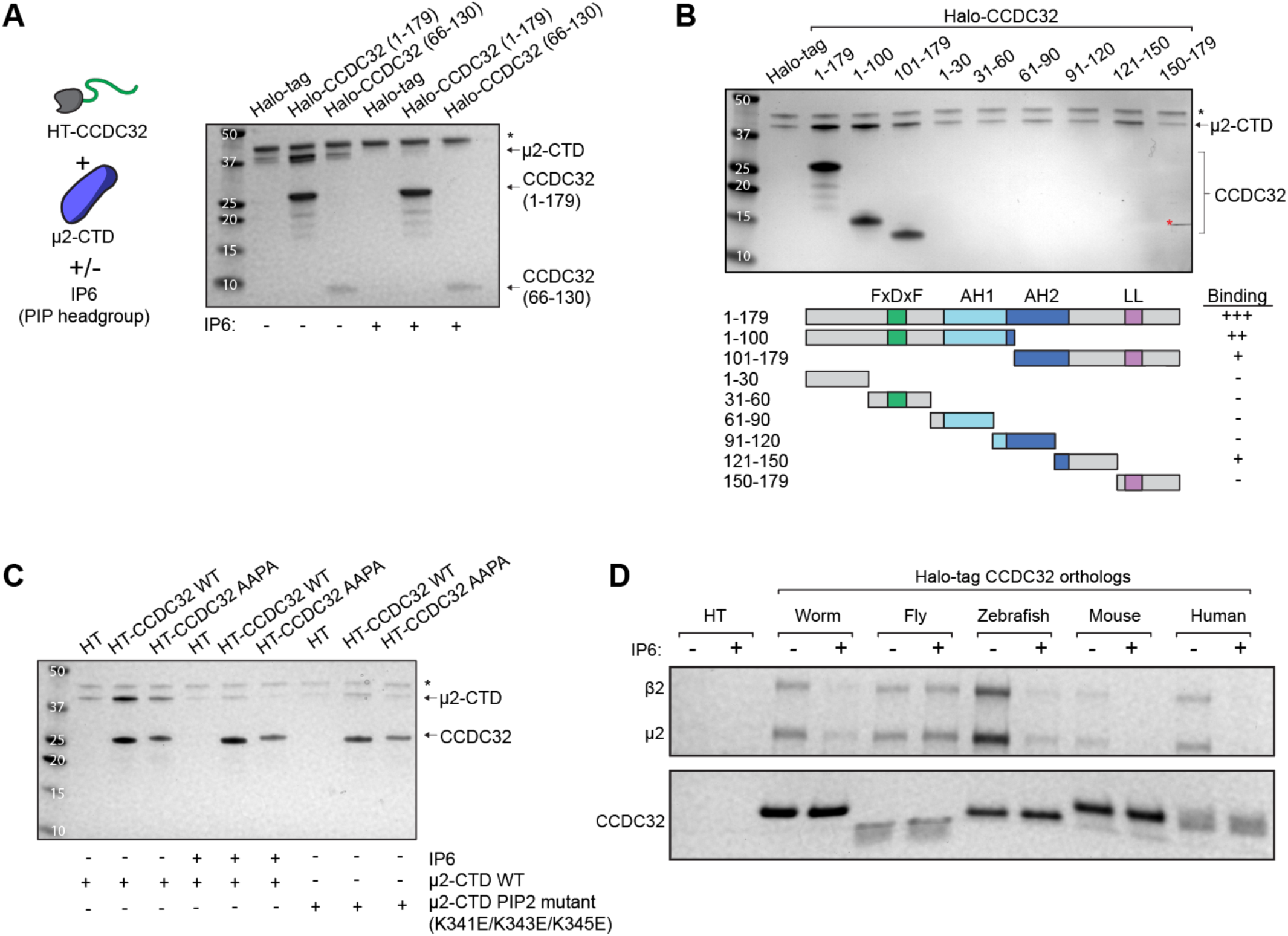
**(E)** Halo-tag pulldown binding assay between various HT-CCDC32 constructs and the μ2-CTD. Addition of the soluble PIP2 headgroup analog IP6 breaks binding. * denotes HRV, used to elute complexes off the Halo resin. **(F)** Halo-tag pulldown binding assay between various HT-CCDC32 truncations and the μ2-CTD. Domain organization is shown below the SDS-PAGE gel, with relative binding strength indicated by -, +, ++, and +++. * as in (**A**). Red * is a contaminant. 30 aa CCDC32 truncations are not visible on the gel. **(G)** Halo-tag pulldown binding assay between HT-CCDC32 constructs and μ2-CTD, either WT or a PIP2-binding mutant bearing a K341E, K343E, K345E triple mutant. Some conditions add IP6, a soluble analog of the PIP2 lipid headgroup. **(H)** Halo-tag pulldown binding assay between HT-CCDC32 ortholog constructs and mouse β2/μ2 hemicomplex. Each ortholog is tested for binding +/-IP6.

**Supplementary Figure 8.**
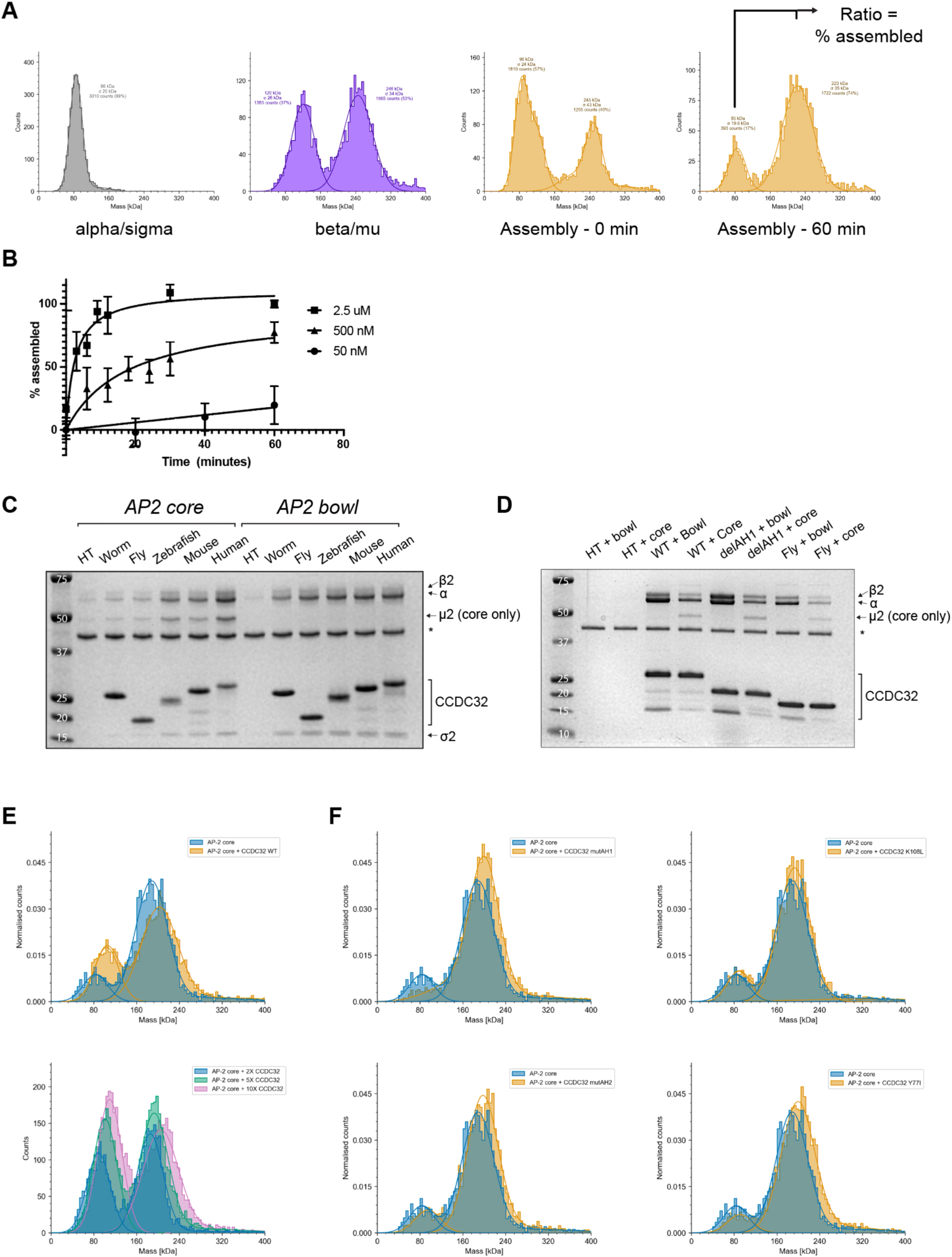
**(A)** Representative mass photometry mass distribution plots for hemicomplexes, assembly at time point 0 min, and assembly at time point 60 minutes. **(B)** Time-course AP-2 assembly assay performed with varying concentrations of hemicomplexes. **(C)** Halo-tag pulldown binding assay between HT-CCDC32 constructs with AP-2 core and AP-2 bowl. Note the enrichment of α vs β2 for all constructs. AP-2 bowl bears a μ2 truncation that removes the μ2-CTD, causing the μ2 NTD to co-migrate with σ2. **(D)** Halo-tag pulldown binding assay between HT-CCDC32 mutant constructs with AP-2 core and AP-2 bowl. Note the ratio of α vs β2. **(E)** Mass photometry of AP-2 alone, or co-incubated with WT CCDC32 (top) or with varying molar ratios of WT CCDC32 (bottom) **(F)** Mass photometry of AP-2 alone, or co-incubated with an equimolar amount of various CCDC32 mutants (mutAH1, mutAH1, K108L, Y77I).

**Supplementary Figure 9.**
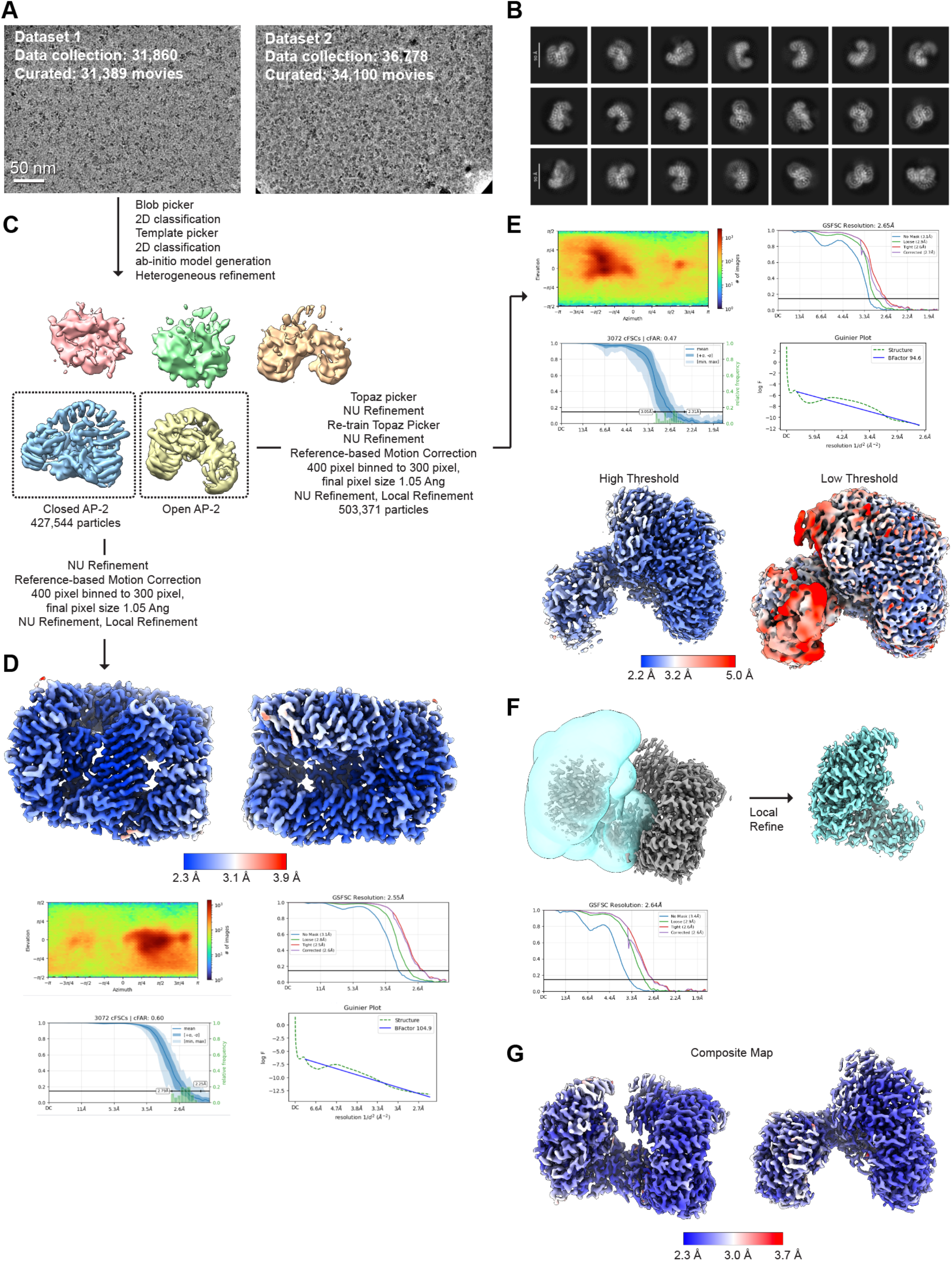
**(A)** Representative images of the two datasets. Both were collected with identical microscope parameters on separate cryo-EM grid samples. **(B)** Representative 2D class averages after multiple rounds of picking and 2D classification. **(C)** Each dataset was processed in a similar manner. First, micrographs were picked with blob picker, then extensively 2D classified. 2D classes from “clean” particles were used as templates for picking, followed by 2D classification and ab-initio model generation with the “clean” particles. Ab-initio classes were used for heterogeneous refinement. The heterogeneous refinement shown here is only for Dataset 1, showing two high-resolution classes. One close corresponds to closed AP-2 with no density for CCDC32 bound. These particles were further processed as outlined. **(D)** Final refinement for closed AP-2. Map is colored by local resolution and refinement statistic plots from cryoSPARC are shown. **(E)** A second high-resolution class from Dataset 1 in panel (**C**) corresponds to open AP-2 bound to the dileucine motif of CCDC32. These particles were further refined as outlined. The consensus refinement is shown with the map colored by local resolution and refinement statistic plots from cryoSPARC included. Note that one lobe of the complex is conformationally heterogeneous and has low local resolution. **(F)** To increase the resolution of the α/σ2 region of the map, a local refinement was performed following the consensus refinement. The refinement mask is shown over the consensus refinement, and the final local refinement is shown in cyan. **(G)** The consensus and local refinement half maps were merged using phenix.combine_focused_maps and used to create a sharpened composite map. The composite map is shown colored by local resolution.

**Supplementary Figure 10.**
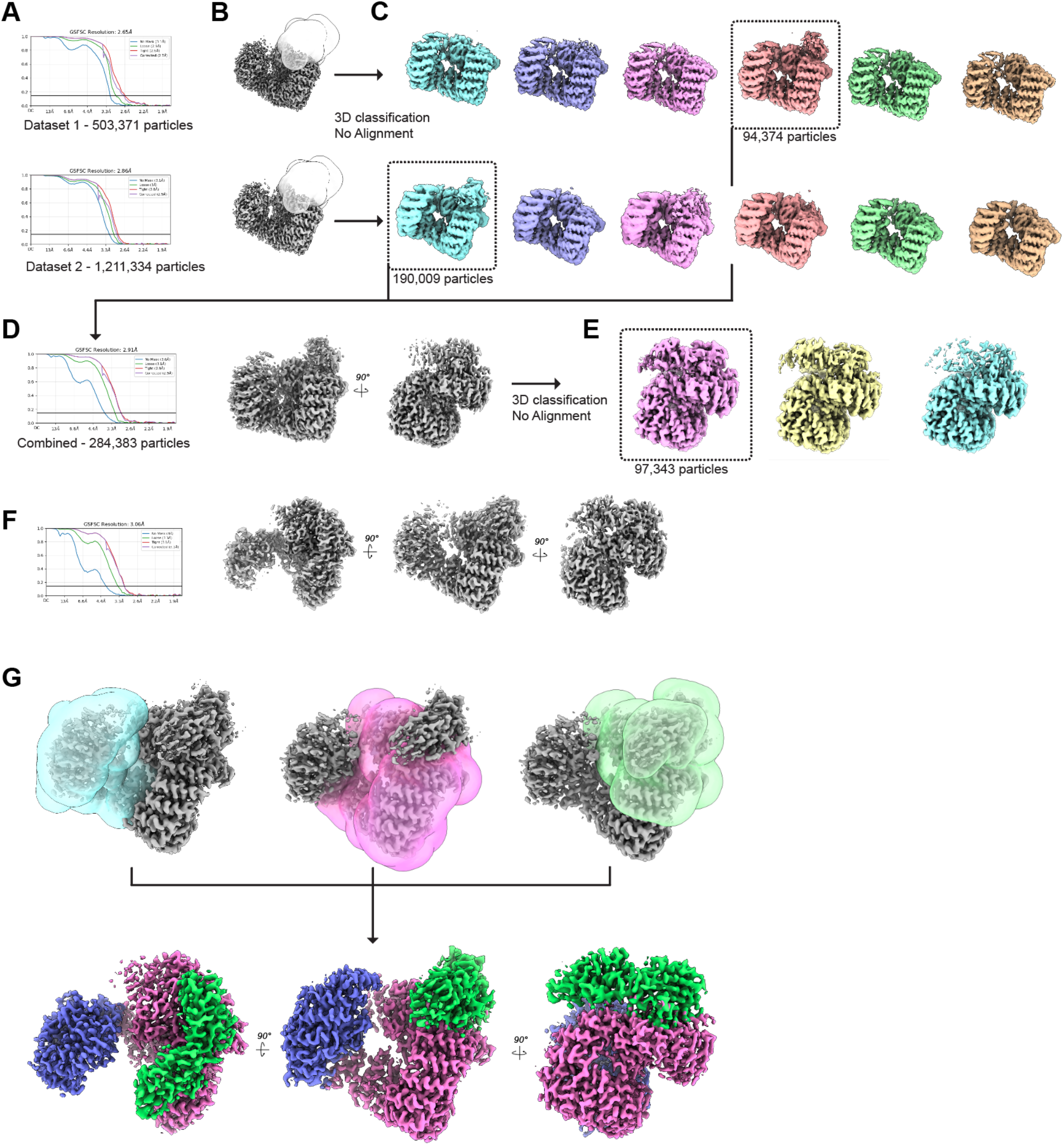
**(A)** Each Dataset was processed independently as outline in Supp. Fig. 8, yielding a set of open AP-2 particles with density for the dileucine motif of CCDC32. Each particle dataset was refined separately against the same 3D reference. **(B)** Focus masks for each dataset are shown for the open AP-2 maps from (**A**). The mask is placed where fragmented density for the μ2-CTD was observed in a subset of 3D classes during processing. **(C)** Each dataset was used for focused 3D classification without alignment. This revealed a 10-15% population of each dataset with a fixed μ2-CTD, revealing a complex in the same general conformation as AP-2 bound to cargo in a clathrin-coated vesicle. **(D)** The cargo-bound AP-2 particles from both datasets were combined and refined. This yielded a complex with better, but still slightly fragmented density for the μ2-CTD, likely do to small changes in the position of the domain. **(E)** A second focused 3D classification without alignment was used to identify 97,343 particles with a rigid conformation of the μ2-CTD. **(F)** The consensus refinement for the cargo-bound AP-2 particles is shown. **(G)** To improve the resolution, three focus refinements were performed. A composite map was made from the half maps using phenix.combine_focused_maps. The contribution from each focus refinement is colored separately.

**Supplementary Figure 11.**
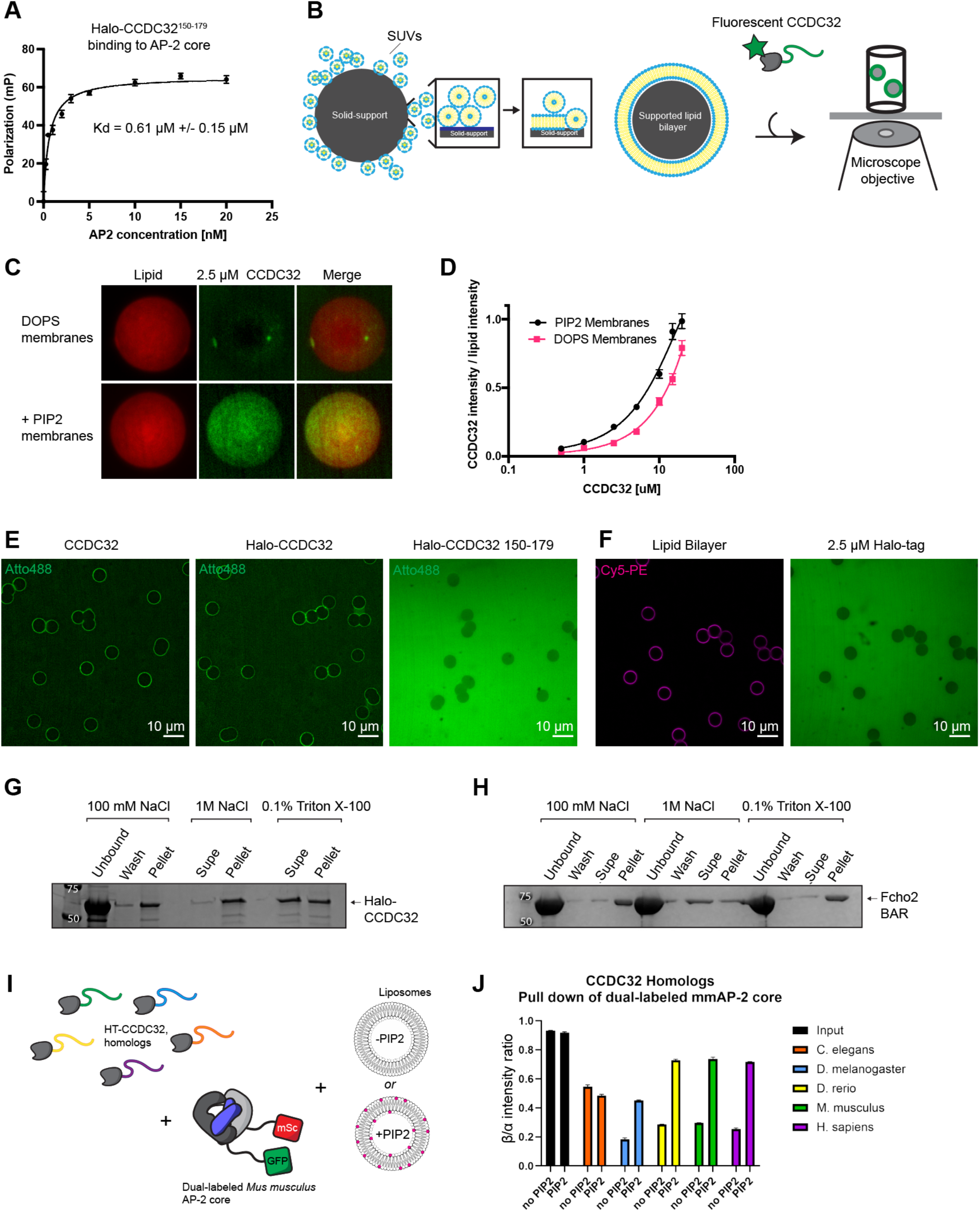
**(A)** Fluorescence Polarization binding assay between Atto488-labeled Halo-CCDC32150-179 and unlabeled AP-2 core. Dissociation constant was determined using linear regression and a single-site binding model. **(B)**Schematic of membrane-binding assay using supported lipid bilayers. Small-unilamellar vesicles (SUVs) of a defined lipid composition are mixed with a solid support, in this case silica microspheres. SUVs spontaneously fuse and form a uniform bilayer on the solid support. SLBs are mixed with fluorescently-tagged protein and imaged with confocal fluorescence microscopy. **(C)** Membrane binding assay between Halo-CCDC32 and membranes containing DOPC/DOPS or DOPC/DOPS + 5% PIP2. Representative SLBs are shown for both conditions, as well as the lipid bilayer (Cy5-PE). **(D)** Binding plot showing incubation of Halo-CCDC32 at various concentrations to both membrane conditions. For each condition, at least 40 SLBs were quantified to obtain the protein/lipid ratio. **(E)** Membrane binding assay using tagless CCDC32, Halo-CCDC32, and Halo-CCDC32^150-159^. Scale bar = 10 microns. **(F)** Membrane binding assay control showing Halo-tag alone does not bind to membranes. Imaging of Cy5-PE (left) shows that the membrane is intact on the solid support. **(G)** SLB pelleting assay showing CCDC32 is washed away in detergent-containing buffer, but not high-salt buffer. **(H)** SLB pelleting assay showing Fcho2 BAR domain is washed away by high-salt buffer, but is unaffected by detergent-containing buffer. **(I)** Schematic of dual-labeled fluorescent AP-2 pull-down assay in the presence of PC/PS liposomes of PC/PS + 5% PIP2 liposomes. **(J)** Bar graph of β2/α fluorescence intensity ratios after a Halo-tag pulldown. All readings are normalized to an input of β2/α. Note: this graph does not show differences in affinity between the CCDC32 constructs, only the ratio of β2/α.

## References

1. Havugimana, P. C. et al. A Census of Human Soluble Protein Complexes. Cell 150, 1068–1081 (2012).

2. Ellis, R. J. Assembly chaperones: a perspective. Philos. Trans. R. Soc. Lond. B. Biol. Sci. 368, 20110398 (2013).

3. Hartl, F. U., Bracher, A. & Hayer-Hartl, M. Molecular chaperones in protein folding and proteostasis. Nature 475, 324–332 (2011).

4. Wilson, R. H. & Hayer-Hartl, M. Complex Chaperone Dependence of Rubisco Biogenesis. Biochemistry 57, 3210–3216 (2018).

5. Quinlan, R. A. & Ellis, R. J. Chaperones: needed for both the good times and the bad times. Philos. Trans. R. Soc. Lond. B. Biol. Sci. 368, 20130091 (2013).

6. Zhang, Q., Yu, D., Seo, S., Stone, E. M. & Sheffield, V. C. Intrinsic Protein-Protein Interaction-mediated and Chaperonin-assisted Sequential Assembly of Stable Bardet-Biedl Syndrome Protein Complex, the BBSome*. J. Biol. Chem. 287, 20625–20635 (2012).

7. Burgess, R. J. & Zhang, Z. Histone chaperones in nucleosome assembly and human disease. Nat. Struct. Mol. Biol. 20, 14–22 (2013).

8. Gulbranson, D. R. et al. AAGAB Controls AP2 Adaptor Assembly in Clathrin-Mediated Endocytosis. Dev. Cell 50, 436–446.e5 (2019).

9. Wan, C. et al. An AAGAB-to-CCDC32 handover mechanism controls the assembly of the AP2 adaptor complex. Proc. Natl. Acad. Sci. U. S. A. 121, e2409341121 (2024).

10. Traub, L. M. & Bonifacino, J. S. Cargo Recognition in Clathrin-Mediated Endocytosis. Cold Spring Harb. Perspect. Biol. 5, (2013).

11. Pearse, B. M. & Robinson, M. S. Purification and properties of 100-kd proteins from coated vesicles and their reconstitution with clathrin. EMBO J. 3, 1951–1957 (1984).

12. Collins, B. M., McCoy, A. J., Kent, H. M., Evans, P. R. & Owen, D. J. Molecular Architecture and Functional Model of the Endocytic AP2 Complex. Cell 109, 523–535 (2002).

13. Ohno, H. et al. Interaction of tyrosine-based sorting signals with clathrin-associated proteins. Science 269, 1872–1875 (1995).

14. Owen, D. J. & Evans, P. R. A structural explanation for the recognition of tyrosine-based endocytotic signals. Science 282, 1327–1332 (1998).

15. Doray, B., Lee, I., Knisely, J., Bu, G. & Kornfeld, S. The gamma/sigma1 and alpha/sigma2 hemicomplexes of clathrin adaptors AP-1 and AP-2 harbor the dileucine recognition site. Mol. Biol. Cell 18, 1887–1896 (2007).

16. Kelly, B. T. et al. A structural explanation for the binding of endocytic dileucine motifs by the AP2 complex. Nature 456, 976 (2008).

17. Kovtun, O., Dickson, V. K., Kelly, B. T., Owen, D. J. & Briggs, J. A. G. Architecture of the AP2/clathrin coat on the membranes of clathrincoated vesicles. Sci. Adv. 6, eaba8381 (2020).

18. Paraan, M. et al. The structures of natively assembled clathrin-coated vesicles. Sci. Adv. 6, eaba8397 (2020).

19. Kelly, B. T. et al. AP2 controls clathrin polymerization with a membrane-activated switch. Science 345, 459–463 (2014).

20. Jackson, L. P. et al. A Large-Scale Conformational Change Couples Membrane Recruitment to Cargo Binding in the AP2 Clathrin Adaptor Complex. Cell 141, 1220–1229 (2010).

21. Kovtun, O. et al. Structure of the membrane-assembled retromer coat determined by cryo-electron tomography. Nature 561, 561–564 (2018).

22. Olesen, L. E. et al. Solitary and Repetitive Binding Motifs for the AP2 Complex α-Appendage in Amphiphysin and Other Accessory Proteins*. J. Biol. Chem. 283, 5099–5109 (2008).

23. Beacham, G. M., Partlow, E. A. & Hollopeter, G. Conformational regulation of AP1 and AP2 clathrin adaptor complexes. Traffic 20, 741–751 (2019).

24. Giehl, K. A. et al. Nonsense mutations in AAGAB cause punctate palmoplantar keratoderma type Buschke-Fischer-Brauer. Am. J. Hum. Genet. 91, 754–759 (2012).

25. Pohler, E. et al. Haploinsufficiency for AAGAB causes clinically heterogeneous forms of punctate palmoplantar keratoderma. Nat. Genet. 44, 1272–1276 (2012).

26. Mattera, R., De Pace, R. & Bonifacino, J. S. The adaptor protein chaperone AAGAB stabilizes AP-4 complex subunits. Mol. Biol. Cell 33, ar109 (2022).

27. Wan, C. et al. AAGAB is an assembly chaperone regulating AP1 and AP2 clathrin adaptors. J. Cell Sci. 134, jcs258587 (2021).

28. Harel, T. et al. Loss of function mutations in CCDC32 cause a congenital syndrome characterized by craniofacial, cardiac and neurodevelopmental anomalies. Hum. Mol. Genet. 29, 1489–1497 (2020).

29. Abdalla, E., Alawi, M., Meinecke, P., Kutsche, K. & Harms, F. L. Cardiofacioneurodevelopmental syndrome: Report of a novel patient and expansion of the phenotype. Am. J. Med. Genet. A. 188, 2448–2453 (2022).

30. Fernandes da Rocha, D., Quental, R., Grangeia, A. & Pinto Moura, C. A novel homozygous deletion in CCDC32 gene causing cardiofacio-neurodevelopmental syndrome: the fourth patient reported. Clin. Dysmorphol. 33, 114–117 (2024).

31. Yang, Z. et al. CCDC32 stabilizes clathrin-coated pits and drives their invagination. BioRxiv Prepr. Serv. Biol. 2024.06.26.600785 (2025) doi:10.1101/2024.06.26.600785.

32. Wainberg, M. et al. A genome-wide atlas of co-essential modules assigns function to uncharacterized genes. Nat. Genet. 53, 638–649 (2021).

33. Jumper, J. et al. Highly accurate protein structure prediction with AlphaFold. Nature 596, 583–589 (2021).

34. Lotthammer, J. M. et al. Metapredict enables accurate disorder prediction across the Tree of Life. 2024.11.05.622168 Preprint at 10.1101/2024.11.05.622168 (2024).

35. Yariv, B. et al. Using evolutionary data to make sense of macromolecules with a ‘face-lifted’ ConSurf. Protein Sci. Publ. Protein Soc. 32, e4582 (2023).

36. Zaccai, N. R. et al. FCHO controls AP2’s initiating role in endocytosis through a PtdIns(4,5)P2-dependent switch. Sci. Adv. 8, eabn2018 (2022).

37. Praefcke, G. J. et al. Evolving nature of the AP2 α-appendage hub during clathrin-coated vesicle endocytosis. EMBO J. 23, 4371–4383 (2004).

38. Abramson, J. et al. Accurate structure prediction of biomolecular interactions with AlphaFold 3. Nature 630, 493–500 (2024).

39. Greenberg, M., DeTulleo, L., Rapoport, I., Skowronski, J. & Kirchhausen, T. A dileucine motif in HIV-1 Nef is essential for sorting into clathrin-coated pits and for downregulation of CD4. Curr. Biol. CB 8, 1239–1242 (1998).

40. Partlow, E. A., Cannon, K. S., Hollopeter, G. & Baker, R. W. Structural basis of an endocytic checkpoint that primes the AP2 clathrin adaptor for cargo internalization. Nat. Struct. Mol. Biol. 29, 339–347 (2022).

41. Le Chatelier, H. L. Sur un énoncé général des lois des équilibres chimiques. Comptes Rendus Hebd. Séances L’Académie Sci. 99, 786–789 (1884).

42. Cannon, K. S., Sarsam, R. D., Tedamrongwanish, T., Zhang, K. & Baker, R. W. Lipid nanodiscs as a template for high-resolution cryo-EM structures of peripheral membrane proteins. 2023.03.07.531120 Preprint at 10.1101/2023.03.07.531120 (2023).

43. Wernick, N. L. B., Haucke, V. & Simister, N. E. Recognition of the tryptophan-based endocytosis signal in the neonatal Fc Receptor by the mu subunit of adaptor protein-2. J. Biol. Chem. 280, 7309–7316 (2005).

44. Giménez-Andrés, M., Copic, A. & Antonny, B. The Many Faces of Amphipathic Helices. Biomolecules 8, 45 (2018).

45. Smith, S. M., Baker, M., Halebian, M. & Smith, C. J. Weak Molecular Interactions in Clathrin-Mediated Endocytosis. Front. Mol. Biosci. 4, 72 (2017).

46. Naudi-Fabra, S. et al. An extended interaction site determines binding between AP180 and AP2 in clathrin mediated endocytosis. Nat. Commun. 15, 5884 (2024).

47. Kaksonen, M. & Roux, A. Mechanisms of clathrin-mediated endocytosis. Nat. Rev. Mol. Cell Biol. 19, 313–326 (2018).

48. Gu, M. et al. AP2 hemicomplexes contribute independently to synaptic vesicle endocytosis. eLife 2, e00190 (2013).

49. Hirst, J. et al. Auxilin depletion causes self-assembly of clathrin into membraneless cages in vivo. Traffic Cph. Den. 9, 1354–1371 (2008).

50. Li, W., Puertollano, R., Bonifacino, J. S., Overbeek, P. A. & Everett, E. T. Disruption of the Murine Ap2β1 Gene Causes Nonsyndromic Cleft Palate. Cleft Palate Craniofacial J. 47, 566–573 (2010).

51. Altschul, S. F. et al. Gapped BLAST and PSI-BLAST: a new generation of protein database search programs. Nucleic Acids Res. 25, 3389–3402 (1997).

52. Begley, M., Aragon, M. & Baker, R. W. A structure-based mechanism for initiation of AP-3 coated vesicle formation. 2024.06.05.597630 Preprint at 10.1101/2024.06.05.597630 (2024).

53. Partlow, E. A. et al. A structural mechanism for phosphorylation-dependent inactivation of the AP2 complex. eLife 8, e50003 (2019).

54. Schindelin, J. et al. Fiji: an open-source platform for biological-image analysis. Nat. Methods 9, 676–682 (2012).

55. Delaglio, F. et al. NMRPipe: a multidimensional spectral processing system based on UNIX pipes. J. Biomol. NMR 6, 277–293 (1995).

56. Lee, W., Tonelli, M. & Markley, J. L. NMRFAM-SPARKY: enhanced software for biomolecular NMR spectroscopy. Bioinforma. Oxf. Engl. 31, 1325–1327 (2015).

57. Maciejewski, M. W. et al. NMRbox: A Resource for Biomolecular NMR Computation. Biophys. J. 112, 1529–1534 (2017).

58. Bruker. APEX2, SAINT, and SADABS. Bruker AXS Inc., Madison, Wisconsin, USA. (2009).

59. McCoy, A. J. et al. Phaser crystallographic software. J. Appl. Crystallogr. 40, 658–674 (2007).

60. Liebschner, D. et al. Macromolecular structure determination using X-rays, neutrons and electrons: recent developments in Phenix. Acta Crystallogr. Sect. Struct. Biol. 75, 861–877 (2019).

61. Afonine, P. V. et al. Towards automated crystallographic structure refinement with phenix.refine. Acta Crystallogr. D Biol. Crystallogr. 68, 352–367 (2012).

62. Emsley, P., Lohkamp, B., Scott, W. G. & Cowtan, K. Features and development of Coot. Acta Crystallogr. D Biol. Crystallogr. 66, 486–501 (2010).

63. Williams, C. J. et al. MolProbity: More and better reference data for improved all-atom structure validation. Protein Sci. Publ. Protein Soc. 27, 293–315 (2018).

64. Mastronarde, D. N. Automated electron microscope tomography using robust prediction of specimen movements. J. Struct. Biol. 152, 36–51 (2005).

65. Afonine, P. V. et al. Real-space refinement in PHENIX for cryo-EM and crystallography. Acta Crystallogr. Sect. Struct. Biol. 74, 531–544 (2018).

